# Structural brain correlates of non-verbal cognitive ability in 5-year-old children: findings from the FinnBrain Birth Cohort study

**DOI:** 10.1101/2023.02.22.529110

**Authors:** Elmo P. Pulli, Saara Nolvi, Eeva Eskola, Elisabeth Nordenswan, Eeva Holmberg, Anni Copeland, Venla Kumpulainen, Eero Silver, Harri Merisaari, Jani Saunavaara, Riitta Parkkola, Tuire Lähdesmäki, Ekaterina Saukko, Eeva-Leena Kataja, Riikka Korja, Linnea Karlsson, Hasse Karlsson, Jetro J. Tuulari

## Abstract

Non-verbal cognitive ability predicts multiple important life outcomes, e.g., school and job performance. It has been associated with parieto–frontal cortical anatomy in prior studies in adult and adolescent populations, while young children have received relatively little attention. We explored the associations between cortical anatomy and non-verbal cognitive ability in 165 5-year-old participants (mean scan age 5.40 years, SD 0.13; 90 males) from the FinnBrain Birth Cohort study. T1-weighted brain magnetic resonance images were processed using FreeSurfer. Non-verbal cognitive ability was measured using the Performance Intelligence Quotient (PIQ) estimated from the Block Design and Matrix Reasoning subtests from the Wechsler Preschool And Primary Scale Of Intelligence (WPPSI-III). In vertex-wise general linear models, PIQ scores associated positively with volumes in left caudal middle frontal and right pericalcarine regions, as well as surface area in left caudal middle frontal, left inferior temporal, and right lingual regions. There were no associations between PIQ and cortical thickness. To the best of our knowledge, this is the first study to examine structural correlates of non-verbal cognitive ability in a large sample of typically developing 5-year-olds. The findings are generally in line with prior findings from older age groups.

## Introduction

Cognitive ability is an important predictor for many important life outcomes (Plomin & Von Stumm, 2018), such as school and academic performance (Deary et al., 2007; Neisser et al., 1996; Strenze, 2007), educational attainment (M. I. Brown et al., 2021), occupational status (M. I. Brown et al., 2021; Lang & Kell, 2020; Schmidt & Hunter, 2004; Strenze, 2007), job performance (Bertua et al., 2005; Neisser et al., 1996; Schmidt & Hunter, 2004, 1998; N. Schmitt, 2014), income (M. I. Brown et al., 2021; Furnham & Cheng, 2017; Lang & Kell, 2020; Neisser et al., 1996), life expectancy (Batty et al., 2007; Whalley & Deary, 2001), and other psychiatric and somatic health outcomes (e.g., alcohol use, see Batty et al., 2006; and obesity, see Chandola et al., 2006). Cognitive ability is considered stable (Deary et al., 2013; Gow et al., 2011) and highly genetically determined (Deary et al., 2006; Plomin & Von Stumm, 2018) individual characteristic in adult populations, while environmental factors play a greater role the younger the subjects are (Haworth et al., 2009; Plomin et al., 1997; Plomin & Von Stumm, 2018). General cognitive ability can be divided into verbal and non-verbal ability. Based on current evidence in school-age children and adolescents, verbal ability is associated with structural and functional neural features in language areas (Khundrakpam et al., 2017; Qi et al., 2019; Ramsden et al., 2011), while non-verbal ability is associated with structural and functional features in (pre)motor areas (Kim et al., 2016; Ramsden et al., 2011). Furthermore, cognitive ability and brain structural features (volume, surface area (SA), and cortical thickness (CT)) are highly heritable in both children (Deary et al., 2006; Jha et al., 2018; Lenroot et al., 2009; J. E. Schmitt et al., 2019; Wallace et al., 2006) and adults (Deary et al., 2006; Panizzon et al., 2009; Posthuma et al., 2002; J. E. Schmitt et al., 2019; Thompson et al., 2001; Winkler et al., 2010).

The developmental research on brain–cognitive correlates is challenged by the dynamic development of brain across the increasing age. The brain grows rapidly in the first years of life, reaching ca. 80% of adult volume by the age 2 years (Knickmeyer et al., 2008), and 95% by the age 6 years (Phan et al., 2018). Total gray matter (GM) volume reaches its peak at approximately 6 years of age (Bethlehem et al., 2022; Courchesne et al., 2000). The development of GM volumes varies depending on the region, wherein frontal and temporal regions show peak volumes in late childhood, while parietal and occipital volumes are already decreasing by the age 5 years (Aubert-Broche et al., 2013). In turn, cortical SA shows global increase in early childhood and reaches its peak at approximately 10 to 12 years of age (Bethlehem et al., 2022; T. T. Brown et al., 2012; Raznahan et al., 2011; Wierenga et al., 2014). There has been controversy regarding the developmental trajectory of CT with estimates of the age of peak CT varying from early to late childhood (Walhovd et al., 2016). However, a recent study combining data from over 100 studies and 100,000 scans has concluded that CT peaks as early as the second year of life (Bethlehem et al., 2022). Notably, some earlier studies have found different developmental trajectories of CT development depending on the child’s cognitive ability (Khundrakpam et al., 2017; Shaw et al., 2006), challenging the idea that simply being further on the typical developmental neural trajectory would correlate with higher cognitive ability. As such, it is important to explore longitudinal samples to characterize the potential individual differences in the developmental trajectories. However, in many previous longitudinal studies on the topic, the follow-up only starts at approximately 5 years of age (Khundrakpam et al., 2017; Shaw et al., 2006; Sowell et al., 2004), losing statistical power in the youngest age groups. Therefore, large cross-sectional samples can be especially useful to provide new information in the less explored young age groups.

The Parieto–Frontal Integration Theory (P–FIT) model proposes that cognitive ability is consistently associated with structural and functional features of a network including widespread frontal and parietal regions, the anterior cingulate cortex, and sensory regions within the temporal and occipital lobes (based on a review of the literature, see Jung & Haier, 2007). A more recent meta-analysis of structural and functional neuroimaging studies found generally good agreement with the P–FIT model, however there was discrepancy in the results regarding the temporal and occipital regions, for example, related to task vs. resting state functional imaging (Basten et al., 2015). However, these findings are mostly based on adult studies (Basten et al., 2015 excluded studies in children and adolescents from their review), while the neural bases of cognitive ability at different ages and developmental stages throughout childhood are not as well understood.

In line with the P-FIT model, previous studies on school-age children and adolescents have found positive associations between general cognitive ability and GM volume in frontal (Pangelinan et al., 2011; Reiss et al., 1996) and parietal lobes (Pangelinan et al., 2011). One study found prefrontal cortical GM volume to predict approximately 20% of the variance in cognitive ability (greater volume predicted higher cognitive ability) in children between the ages 5 and 17 years (Reiss et al., 1996). Additionally, studies of children and adolescents have found negative associations between general cognitive ability and the volumes in the right middle temporal gyrus (Yokota et al., 2015 participants separated into clusters with different profiles of cognitive ability) as well as positive associations between general cognitive ability and GM volumes in the whole brain and the bilateral cingulate gyrus (Wilke et al., 2003, effects were driven by the adolescents). There is some evidence that SA is also positively associated with general cognitive ability from birth to 11 years of age (Girault et al., 2020; Schnack et al., 2015; Sølsnes et al., 2015) and that children with higher cognitive ability reach the maximal SA faster (Schnack et al., 2015). Furthermore, greater prefrontal SA has been linked to higher general cognitive ability in children aged 9 to 11 years (Vargas et al., 2020). However, pediatric studies examining the connection between SA and cognitive ability are scarce relative to CT studies.

Also, in line with the P-FIT model, greater CT in frontal and parietal regions may predict later higher verbal ability in infants (Girault et al., 2020) or academic achievement in adolescents (Meruelo et al., 2019). Similarly, studies have found positive associations between non-verbal ability and CT in frontal regions in 4– 7-year-old children (with low socioeconomic status, please see Leonard et al., 2019) and adolescents (Schilling et al., 2013). On the other hand, a study of 12–14-year-olds found negative associations between general cognitive ability and CT in bilateral parietal regions (Squeglia et al., 2013). Similarly, one study found negative associations between CT and working memory in 4–8-year-olds in superior and middle frontal, superior parietal, and anterior cingulate regions (Botdorf & Riggins, 2018), while another found no correlations between CT and working memory in any brain regions in 6–16-year-olds (Faridi et al., 2015). Furthermore, a recent longitudinal study in children and adolescents found positive correlations between general cognitive ability and CT mostly in the superior frontoparietal cortex, frontopolar cortex, and language centers (J. E. Schmitt et al., 2019), which are among the areas typically associated with cognitive ability according to the P-FIT model (Jung & Haier, 2007). Notably, correlations were modest in young children but became stronger at approximately 10 years of age. Some other studies have also focused on this dynamic development of CT in childhood and adolescence: One study found greater vocabulary improvement associated with greater thinning between the ages 5 and 11 years in widespread brain regions especially in the left hemisphere (Sowell et al., 2004). In another study, the correlation between general cognitive ability and CT was negative until about 8 years of age and then turned positive (Shaw et al., 2006).

In summary, most studies examining brain structure and cognitive ability are conducted in samples with wide age ranges typically focusing on late childhood and adolescence, while such research in younger age groups is scarcer. Notably, studies with wider age ranges risk conflating findings from different age groups, and studies with large samples from a limited age range are warranted to better explore the neural basis of cognitive ability at the specific developmental stage. To the best of our knowledge, there are no previous large neuroimaging studies focusing solely on typically developing 5-year-olds. Five years is a particularly interesting age to study the structural brain correlates of cognitive ability, as the children are old enough to both cooperate in cognitive assessment to be reliably evaluated and to lie still in the scanner while awake. Furthermore, 5-year-olds have yet to start school, meaning most of them haven’t gone through the changes associated with academic abilities such as reading (Chyl et al., 2021) and arithmetic (Hashimoto et al., 2022).

In the current study, we examined cortical structural correlates of non-verbal ability at 5 years of age. More specifically, we explored the associations between cortical gray matter volume, SA, and CT and non-verbal ability measured with Block Design and Matrix Reasoning tasks from the Wechsler Preschool and Primary Scale of Intelligence (WPPSI-III; Wechsler, 1967) in a sample of 165 typically developing 5-year-olds participating in a larger birth cohort follow-up. Based on previous research, we expected non-verbal ability to be positively associated with volume and SA in frontal and parietal regions. We also expected to find associations between cognitive ability and CT in frontal and parietal regions, but we did not set an explicit hypothesis for the direction of the association, as the findings from previous CT studies are conflicting and studies regarding a similar age group are rare.

## Methods

This study was conducted in accordance with the Declaration of Helsinki, and it was approved by the Joint Ethics Committee of the University of Turku and the Hospital District of Southwest Finland: 1) ETMK: 26/1801/2015 for the neuropsychological measurements, and 2) ETMK: 31/180/2011 for the neuroimaging.

### Participants

The participants are a part of the FinnBrain Birth Cohort Study (finnbrain.fi), which prospectively examines the influence of genetic and environmental factors on child development and later health outcomes (Karlsson et al., 2018). Pregnant women (n = 3808) attending their first trimester ultrasound at gestational week (GW) 12, their spouses (n = 2623), and babies to-be born (n = 3837; including 29 twin pairs) were recruited in Southwest Finland between December 2011 and April 2015. Ultrasound-verified pregnancy and a sufficient knowledge of Finnish or Swedish languages were required for participation. The cohort study includes several follow-up studies. Those participants that attended neuropsychological and neuroimaging visits as part of the 5-year data collection were included in this study.

The participants were first recruited to the neuropsychological assessments at 5 years of age. The participants recruited for this visit were focus cohort families (highest or lowest quartile scores of maternal prenatal distress, please see Karlsson et al., 2018 for more details) and families who had actively participated in previous FinnBrain study visits. For the neuroimaging visit, we primarily recruited participants that had attended the neuropsychological visit. For the neuropsychological visits, 1288 families were contacted and informed of the study, and of these families 974 (75.6%) were reached by telephone. From all the contacted families, 545 (42.3%) participated in a study visit (304 boys (55.8%), mean age 5.01 (SD 0.08), range 4.89– 5.37 years). For the neuroimaging visits, 541 families were contacted and 478 (88.4%) of them were reached. In total, 203 (37.5%) participants attended imaging visits (113 boys (55.7%), mean age 5.40 (SD 0.13), range 5.08–5.79 years). Altogether 196 participants attended both visits.

We originally aimed to scan all subjects between the ages 5 years 3 months and 5 years 5 months; however, there was a pause in visits due to the start of the COVID-19 pandemic, and subsequently many of the participants were older than planned when they were scanned (152/203 (74.9%) of the participants attended the visit within the intended age range). The exclusion criteria for the neuroimaging study were: 1) born before GW 35 (before GW 32 for those with exposure to maternal prenatal synthetic glucocorticoid treatment), 2) developmental anomaly or abnormalities in senses or communication (e.g., blindness, deafness, congenital heart disease), 3) known long-term medical diagnosis (e.g., epilepsy, autism), 4) ongoing medical examinations or clinical follow up in a hospital (meaning there has been a referral from primary care setting to special health care), 5) child use of continuous, daily medication (including per oral medications, topical creams and inhalants; One exception to this was desmopressin medication, which was allowed), 6) history of head trauma (defined as concussion necessitating clinical follow up in a health care setting or worse), 7) metallic (golden) ear tubes (to assure good-quality scans), and routine magnetic resonance imaging (MRI) contraindications.

For this study, only participants with an adequate quality T1 image (n = 173/203, assessed by E.P.P. as described in Pulli et al., 2022) and successful assessment of cognition (n = 166/173) were included. Additionally, one participant was excluded due to scoring below 4 scaled score in verbal ability test Similarities and below the standard score 70 assessed by the performance intelligence quotient (PIQ; calculated from Block Design and Matrix Reasoning scaled scores and the estimated scaled score for a third non-verbal subtest, see more detailed description later in the Methods), leaving us with a final sample size of 165 participants. A few participants were missing one of the non-verbal tasks, and missing data was not imputed. Consequently, the sample sizes were 164 for the Matrix Reasoning task, 160 for the Block Design task, and 159 for PIQ. The characteristics of the final sample (n = 165) are displayed in Table.

**Table.**
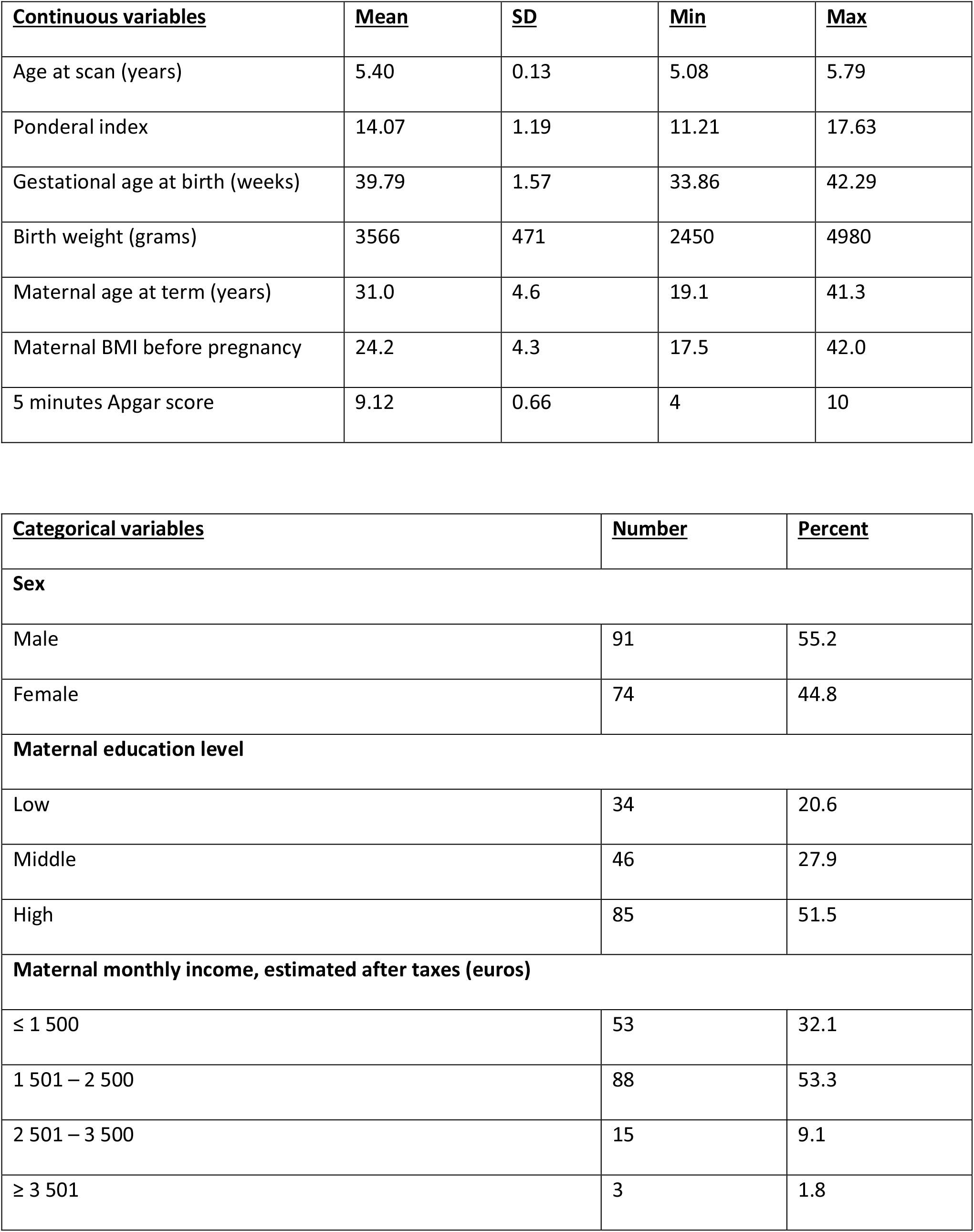
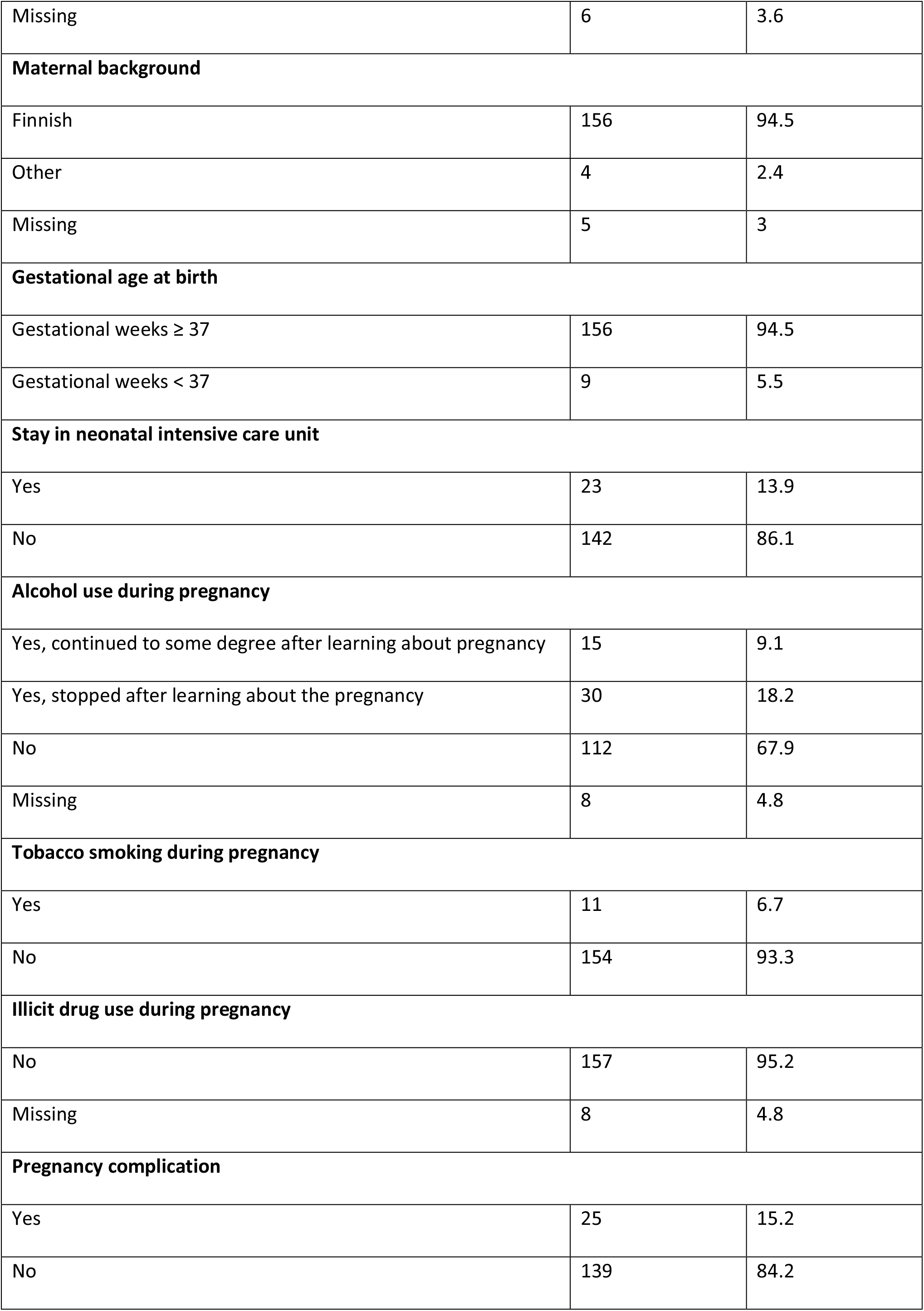
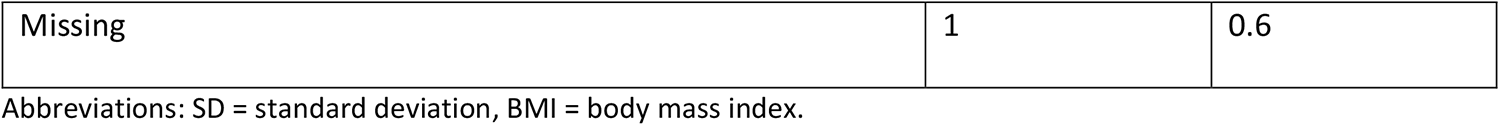
Participant demographics and maternal medical history variables. Number of participants = 165.

| <u>Continuous variables</u> | <u>Mean</u> | <u>SD</u> | <u>Min</u> | <u>Max</u> |
| --- | --- | --- | --- | --- |
| Age at scan (years) | 5.40 | 0.13 | 5.08 | 5.79 |
| Ponderal index | 14.07 | 1.19 | 11.21 | 17.63 |
| Gestational age at birth (weeks) | 39.79 | 1.57 | 33.86 | 42.29 |
| Birth weight (grams) | 3566 | 471 | 2450 | 4980 |
| Maternal age at term (years) | 31.0 | 4.6 | 19.1 | 41.3 |
| Maternal BMI before pregnancy | 24.2 | 4.3 | 17.5 | 42.0 |
| 5 minutes Apgar score | 9.12 | 0.66 | 4 | 10 |

| <u>Categorical variables</u> | <u>Number</u> | <u>Percent</u> |
| --- | --- | --- |
| **Sex** |  |  |
| Male | 91 | 55.2 |
| Female | 74 | 44.8 |
| **Maternal education level** |  |  |
| Low | 34 | 20.6 |
| Middle | 46 | 27.9 |
| High | 85 | 51.5 |
| **Maternal monthly income, estimated after taxes (euros)** |  |  |
| ≤ 1 500 | 53 | 32.1 |
| 1 501 – 2 500 | 88 | 53.3 |
| 2 501 – 3 500 | 15 | 9.1 |
| ≥ 3 501 | 3 | 1.8 |
| Missing | 6 | 3.6 |
| **Maternal background** |  |  |
| Finnish | 156 | 94.5 |
| Other | 4 | 2.4 |
| Missing | 5 | 3 |
| **Gestational age at birth** |  |  |
| Gestational weeks $\geq 37$ | 156 | 94.5 |
| Gestational weeks $< 37$ | 9 | 5.5 |
| **Stay in neonatal intensive care unit** |  |  |
| Yes | 23 | 13.9 |
| No | 142 | 86.1 |
| **Alcohol use during pregnancy** |  |  |
| Yes, continued to some degree after learning about pregnancy | 15 | 9.1 |
| Yes, stopped after learning about the pregnancy | 30 | 18.2 |
| No | 112 | 67.9 |
| Missing | 8 | 4.8 |
| **Tobacco smoking during pregnancy** |  |  |
| Yes | 11 | 6.7 |
| No | 154 | 93.3 |
| **Illicit drug use during pregnancy** |  |  |
| No | 157 | 95.2 |
| Missing | 8 | 4.8 |
| **Pregnancy complication** |  |  |
| Yes | 25 | 15.2 |
| No | 139 | 84.2 |
| Missing | 1 | 0.6 |

Abbreviations: SD = standard deviation, BMI = body mass index.

Ponderal index was calculated using the following formula: weight in kilograms divided by height in meters cubed. Height and weight were acquired during the neuroimaging visit. The participants kept indoor clothes on during the weighing.

Maternal education data was combined from questionnaire data from 14 weeks gestation or 5 years of child age by choosing the highest degree reported. The three classes are: Low = Upper secondary school or vocational school or lower, Middle = University of applied sciences, High = University.

On the question about alcohol usage, four subjects answered that they did not use alcohol during pregnancy, but also answered that they stopped using alcohol when they learned about the pregnancy. These were classified as “yes, stopped when learning about pregnancy”.

The data for maternal monthly income estimate, alcohol use, and illicit drug use are from questionnaires at gestational week 14.

The pregnancy complications include a diagnosis (according to ICD-10) for O12 (Gestational edema and proteinuria without hypertension), O13 (Gestational hypertension without significant proteinuria), O14 (Severe pre-eclampsia), O24 (Diabetes mellitus in pregnancy, childbirth, and the puerperium), O46 (Antepartum hemorrhage, not elsewhere classified), or O99.0 (Anemia complicating pregnancy, childbirth and the puerperium).

Sex, birth weight, maternal BMI before pregnancy, and smoking data (combined with questionnaire data) were retrieved from the National Institute for Health and Welfare (www.thl.fi).

### Bias assessment

Mothers of the children who did not participate in the neuropsychological visits (out of the 1288 contacted families) had a lower education level (χ^2^(2) = 30.94, p < 0.001), a lower monthly income (χ^2^(3) = 11.65, p = 0.009) and were younger (*t* (1286) = -4.130, p < 0.001) compared to the mothers in the families that participated in the neuropsychological visits.

Mothers of the children who participated in the neuropsychological visits but not in the neuroimaging visits were older (*t* (369) = 1.97, p = 0.047) but did not differ in education level or monthly income compared to the mothers in the families that participated in the MRI visit.

The children who participated in the neuropsychological visits but not in the MRI visits did not differ in PIQ, Block Design, or Matrix Reasoning performance from those that participated in the MRI visit.

### Procedures

Non-verbal ability was assessed at 5 years of age using the Block Design and Matrix Reasoning subtasks of the Wechsler Preschool and Primary Scale of Intelligence-Third Edition (WPPSI-III, Wechsler, 1967).

After the FinnBrain Child Development and Parental Functioning Lab visit, the participating families were invited to the MRI visit, where structural T1-weighted images were collected as a part of max. 60-minute scan.

### Neuropsychological study visits

The neuropsychological study visits for 5-year-old children included neurocognitive testing, eye-movement tracking, mother-child interaction assessment, and questionnaires filled out by the parents. Neurocognitive testing included assessments of the child’s general cognitive ability (WPPSI-III subtests Block Design, Matrix Reasoning and Similarities) and executive functioning and self-regulation, of which only the non-verbal tasks from the general cognitive ability assessments are used in the current study.

The approximately two-hour-long study visits were conducted and video recorded by graduate students in quiet examination rooms and the data collection was overseen by PhD students/psychologists. The graduate students were trained by PhD students/psychologists prior to data collection, to ensure unified test administration among all students, and to ensure that the students had sufficient interaction skills to scaffold the children’s motivation and mood during the study visit. Written informed consent was provided by the parents prior to the study visit, and the parents received feedback of the child’s performance on some of the assessment methods after the study visit.

### Non-verbal ability

Non-verbal ability was assessed using the Finnish version of WPPSI-III, which is a standardized and widely used measure of cognitive ability in young children from ages 2 years and 6 months to 7 years and 3 months (Wechsler, 2009). In this study, a composite sum score of non-verbal ability (PIQ; mean 100) was estimated using two subtests: the Block Design task measuring visuospatial ability and the Matrix Reasoning task measuring visual abstract reasoning. The standardized scale scores corresponding the raw scores of the subtests were based on Finnish norms and result in a mean of 10, reflecting standardized mean performance in the population at each age. Additionally, analyses of the subtests were conducted separately to get further information on the possible subtest driving the findings.

The PIQ scores were: mean = 104.7, SD = 15.4, range 68–146. Block Design scores: mean = 10.5, SD = 3.3, range 3–19. Matrix Reasoning scores: mean = 10.8, SD = 2.8, range 1–18, suggesting normally distributed cognitive ability in the sample of the present study. The Pearson correlation between PIQ and Block Design was 0.809 (p < 0.001), between PIQ and Matrix Reasoning was 0.711 (p < 0.001), and between Block Design and Matrix Reasoning was 0.164 (p = 0.039). Scatter plots of the cognitive measurements are shown in Supplementary Figure 1.

### Neuroimaging study visits

All visits were performed by research staff for research purposes. The participants were recruited via phone calls by the research staff. A staff member made a home visit to deliver practice materials, give further information about the study visit, and to answer any remaining questions. At the start of the study visit, written informed consent from both parents as well as verbal assent from the child were acquired. The visits had a two-hour preparation time before the scan, which consisted of familiarization with the research staff, practice for the scan, and a light meal. The preparation time was long enough so that it allowed the staff to attend to the needs of the child. Participants were scanned awake or during natural sleep. A parent and a research staff member were present in the scanning room throughout the scan. Everyone in the room had their hearing protected with earplugs and headphones. During the scan, participants were allowed to watch a movie or a cartoon of their choice, apart from the functional MRI (fMRI) sequence. The study visit protocol has been described in more detail in our earlier work (Copeland et al., 2021; Pulli et al., 2022).

All images were viewed by one neuroradiologist (R.P.) who then consulted a pediatric neurologist (T.L.) when necessary. The protocol with incidental findings has been described in our earlier work (Kumpulainen et al., 2020). In the whole neuroimaging sample (n = 203), there were 13 participants with incidental findings (6.4%). Of them, 11 were included in the sample of this study (n = 165).

### MRI data acquisition

Participants were scanned using a Siemens Magnetom Skyra fit 3T with a 20-element head/neck matrix coil. We used Generalized Autocalibrating Partially Parallel Acquisition (GRAPPA) technique to accelerate image acquisition (parallel acquisition technique [PAT] factor of 2 was used). The max. 60-minute scan protocol included a high resolution T1 magnetization prepared rapid gradient echo (MPRAGE), a T2 turbo spin echo (TSE), a 7-minute resting state functional MRI, and a 96-direction single shell (b = 1000 s/mm^2^) diffusion tensor imaging (DTI) sequence (Merisaari et al., 2019; Rosberg et al., 2022) as well as a 31-direction with b = 650 s/mm^2^ and a 80-direction with b = 2000 s/mm^2^. For the purposes of the current study, we acquired high resolution T1-weighted images with the following sequence parameters: repetition time = 1900 ms, echo time = 3.26 ms, inversion time = 900 ms, flip angle = 9 degrees, voxel size = 1.0 × 1.0 × 1.0 mm^3^, field-of-view 256 × 256 mm^2^. The scans were planned as per recommendations of the FreeSurfer developers (https://surfer.nmr.mgh.harvard.edu/fswiki/FreeSurferWiki?action=AttachFile&do=get&target=FreeSurfer_Suggested_Morphometry_Protocols.pdf, at the time of writing).

### Image processing

The cortical reconstruction and volumetric segmentation for all 165 images were performed with the FreeSurfer software suite, version 6.0.0 (http://surfer.nmr.mgh.harvard.edu/). We selected the T1 image with the least motion artefact (in case there were several attempts due to visible motion during scan) and then applied the “recon-all” processing stream with default parameters. It begins with transformation to Talairach space, intensity inhomogeneity correction, bias field correction (Sled et al., 1998), and skull-stripping (Ségonne et al., 2004). Thereafter, white matter is separated from gray matter and other tissues and the volume within the created gray–white matter boundary is filled. After this, the surface is tessellated and smoothed. After these preprocessing steps are completed, the surface is inflated (Fischl, Sereno, & Dale, 1999) and registered to a spherical atlas. This method adapts to the folding pattern of each individual brain, utilizing consistent folding patterns such as the central sulcus and the sylvian fissure as landmarks, allowing for high localization accuracy (Fischl, Sereno, Tootell, et al., 1999). FreeSurfer uses probabilistic approach based on Markov random fields for automated labeling of brain regions. CT is calculated as the average distance between the gray–white matter boundary and the pial surface on the tessellated surface (Fischl & Dale, 2000). The CT measurement technique has been validated against manual measurements from imaging data (Kuperberg et al., 2003; Salat, 2004) and against post-mortem histological analysis (Rosas et al., 2002).

After the initial FreeSurfer processing, we visually checked and manually edited all images. Briefly, the manual edits included removing skull fragments where they affected the pial border, correcting errors in the border between gray and white matter, and removing arteries. After the edits, the FreeSurfer recon-all was run again. For a more detailed description of the image processing procedure, please see our previous article (Pulli et al., 2022).

### Confounders

Based on previous studies and our own previous work on this age group (Silver et al., 2022), child sex, age at scan, ponderal index (mass in kilograms divided by height in meters cubed; measured during the neuroimaging visit), as well as maternal age at term and maternal education level were included as covariates in our analyses. Maternal education data was combined from questionnaire data from 14 weeks gestation or 5 years of child age by choosing the highest degree reported (three classes: Low = Upper secondary school or vocational school or lower, Middle = University of applied sciences, High = University; low and middle level education grouped together for statistical analyses).

Additionally, some factors that could potentially affect the results were explored in sensitivity analyses: maternal pre-pregnancy body mass index (BMI; Li et al., 2016; Ou et al., 2015), alcohol exposure *in utero* (mothers who continued to use alcohol after they learned about the pregnancy excluded)(Donald et al., 2015), tobacco exposure *in utero* (mothers with any tobacco use during pregnancy excluded; Chang et al., 2016; Knickmeyer et al., 2016), preterm birth (participants born before GW 37 excluded; Jeong et al., 2016; Kapellou et al., 2006), 5 minutes Apgar score (Aoki et al., 2020; Hong & Lee, 2018), pregnancy complications (mothers with any complications excluded; Xuan et al., 2020), and stay in the neonatal intensive care unit (NICU; those with NICU stay excluded; Aoki et al., 2020). There was some missing data. Eight participants were missing data for the alcohol exposure, and one was missing data for pregnancy complication and in these cases, we used mode imputation. One participant was missing pre-pregnancy maternal BMI, and we used mean imputation based on the 173 participants with available maternal BMI and usable MRI data.

### Statistics

Statistical analyses concerning demographics and regions of interest (ROI) were conducted using the IBM SPSS Statistics for Windows, version 27.0 (IBM Corp., Armonk, NY, USA). Scatter plots and the related statistics were created using JASP version 0.16.1.0 (JASP Team, 2022). Statistical significance in all analyses was calculated two-tailed at alpha level 0.05.

To assess potential selection bias, comparisons between the neuroimaging participants that were included in the final sample (n = 165) and those that were excluded (n = 38) were performed with independent samples t-tests for continuous background factors and chi-square tests for categorical factors. All background factors from Table were examined. Information regarding alcohol exposure was missing for 12/203 participants and was analyzed with three different approaches: 1) missing = no exposure, 2) missing = exposure, and 3) missing data not imputed. For the other background factors, there were 0–7 missing observations and missing data was not imputed for this analysis. P < 0.05 was considered significant in this analysis.

#### Vertex-wise statistical analyses

For the purposes this study, we pre-smoothed fsaverage surfaces as instructed by FreeSurfer manual for analyses with Query, Design, Estimate, Contrast (Qdec), a single-binary application included in the FreeSurfer software suite (www.surfer.nmr.mgh.harvard.edu). Qdec is a graphical user interface for a statistics engine running a vertex-by-vertex general linear model (GLM). For display purposes, we used the standard FreeSurfer’s fsaverage in MNI305 space (MNI = Montreal Neurological Institute). We tested for clusters with statistically significant associations between non-verbal ability and cortical GM volume, SA, and CT. The data was smoothed with a kernel of 10 mm full width at half maximum. A Monte Carlo Null-Z Simulation was run with a z-value threshold of 1.3, corresponding to p = 0.05 (Hagler et al., 2006). After the simulation, a z-value threshold 1.3 was used for statistically significant clusters. For confounding factors and performed sensitivity analyses, please see “Confounders”. Age at scan was squared for the purposes of running Qdec. In the sensitivity analyses, we added the potential confounders to the model one at a time (continuous factors) or excluded the exposed group from the analysis (categorical factors).

#### Region of interest-based analyses

Additionally, we calculated partial correlations (controlling for participant sex, maternal education level, maternal age at term, participant ponderal index at scan, and participant age at scan) between 1) all cognitive measurements (PIQ, Block Design, and Matrix Reasoning), and 2) a multitude of brain metrics, including volume, SA, and CT in all 68 ROIs in the Desikan–Killiany atlas (Desikan et al., 2006) as well as total SA (separately for both hemispheres), mean CT (separately for both hemispheres), brain volume (excluding ventricles), and estimated total intracranial volume; in total 210 brain metrics per cognitive measurement. We excluded poor quality ROIs from this analysis (described in detail in our previous article Pulli et al., 2022). For this part of the analysis, we corrected for multiple comparisons using Bonferroni correction (630 comparisons corresponding to Bonferroni-corrected p-value = 0.05/630 = 0.000079).

## Results

### Demographics

The children from the neuroimaging sample (n = 203) that were included in this study (n = 165) had higher gestational age at birth (included 39.79 weeks, SD 1.57; excluded 39.01 weeks, SD 2.15; p = 0.041) and fewer had a NICU stay (included 142 no, 23 yes (13.9%); excluded 25 no, 11 yes (30.6%); χ2(1) = 0.016), and their mothers had lower pre-pregnancy BMI (included 24.20, SD 4.33; excluded 25.83, SD 4.77; p = 0.044), compared to those who were excluded (n = 38).

### Cortical gray matter volume and non-verbal ability

Figure 1 presents the associations between brain volume and non-verbal ability. All significant associations were positive. For PIQ, there were significant clusters in the left caudal middle frontal gyrus (peak z = 1.67, size = 950.9 mm^2^, peak coordinates -41.4, 3.5, 46.8) and the right pericalcarine region (peak z = 4.00, size = 1639.8 mm^2^, peak coordinates 14.3, -77.4, 4.8). There were no significant correlations between Block Design scores and brain volumes. For Matrix Reasoning, there was a significant cluster in the right pericalcarine region (peak z = 4.00, size = 1859.4 mm^2^, peak coordinates 14.3, -77.4, 4.8).

**Figure 1.**
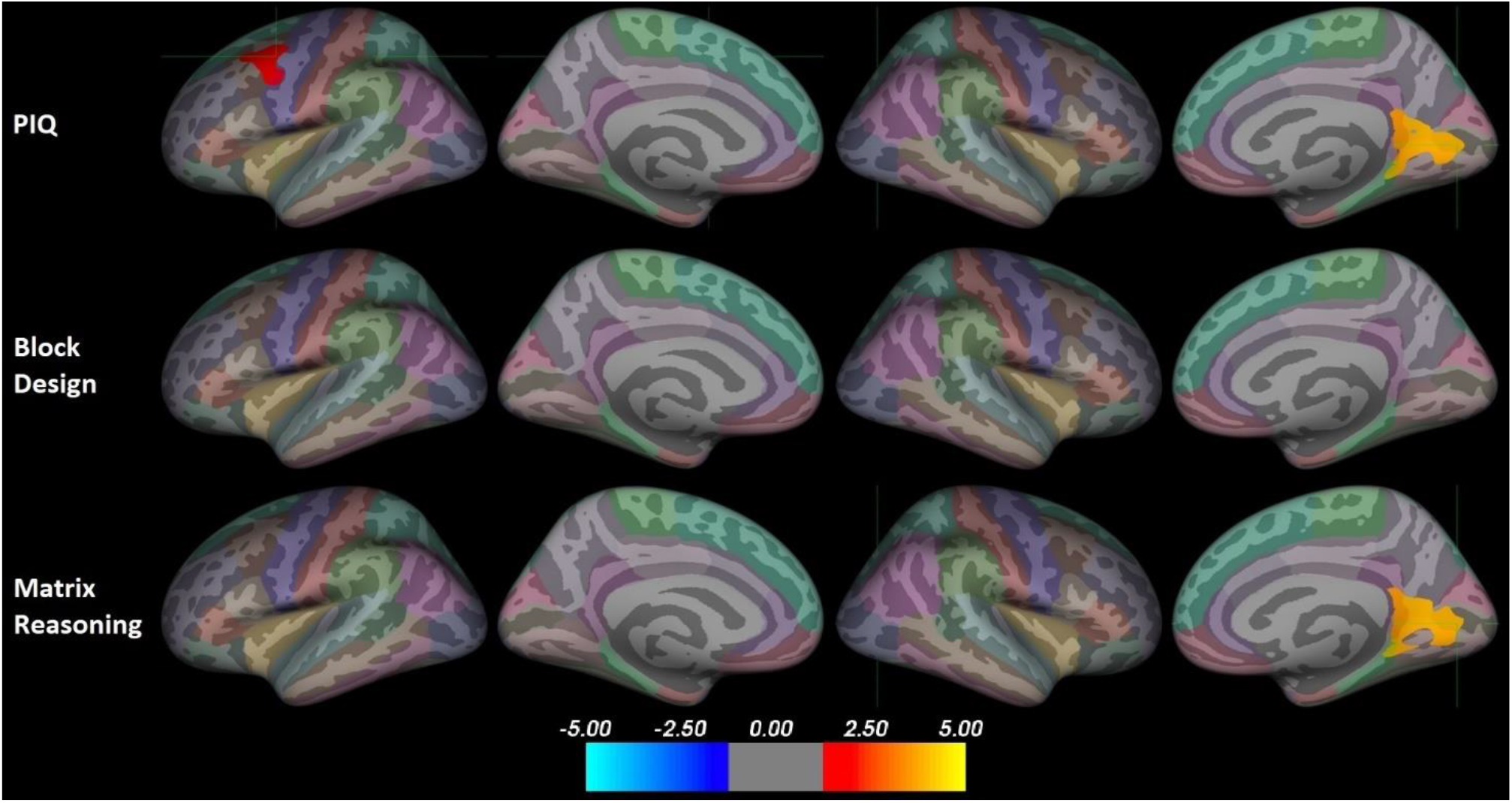
Positive associations between non-verbal ability and cortical gray matter volume. Results corrected for multiple comparisons using Monte-Carlo Null-Z simulation. Color indicates significance as a z-value. The position of the green crosshair indicates the most statistically significant vertex in statistically significant clusters. Left hemisphere on the left and right on the right side. Color coding of regions according to the Desikan–Killiany atlas. PIQ = performance intelligence quotient.

### Pial surface area and non-verbal ability

Figure 2 presents the associations between brain SA and non-verbal ability. All significant associations were positive. For PIQ, there were significant clusters in the left caudal middle frontal gyrus (peak z = 2.05, size = 870.7 mm^2^, peak coordinates -36.1, 0.6, 31.0), the left inferior temporal gyrus (peak z = 1.37, size = 692.4 mm^2^, peak coordinates -45.0, -18.5, -29.5), and the right lingual gyrus (peak z = 3.70, size = 1239.1 mm^2^, peak coordinates 25.4, -61.6, 0.7). There were no significant correlations between Block Design scores and SA. For Matrix Reasoning, there were significant clusters in the left caudal middle frontal gyrus (peak z = 2.07, size = 874.9 mm^2^, peak coordinates -36.1, 0.6, 31.0) and the right lingual gyrus (peak z = 4.00, size = 1311.8 mm^2^, peak coordinates 25.4, -61.6, 0.7).

**Figure 2.**
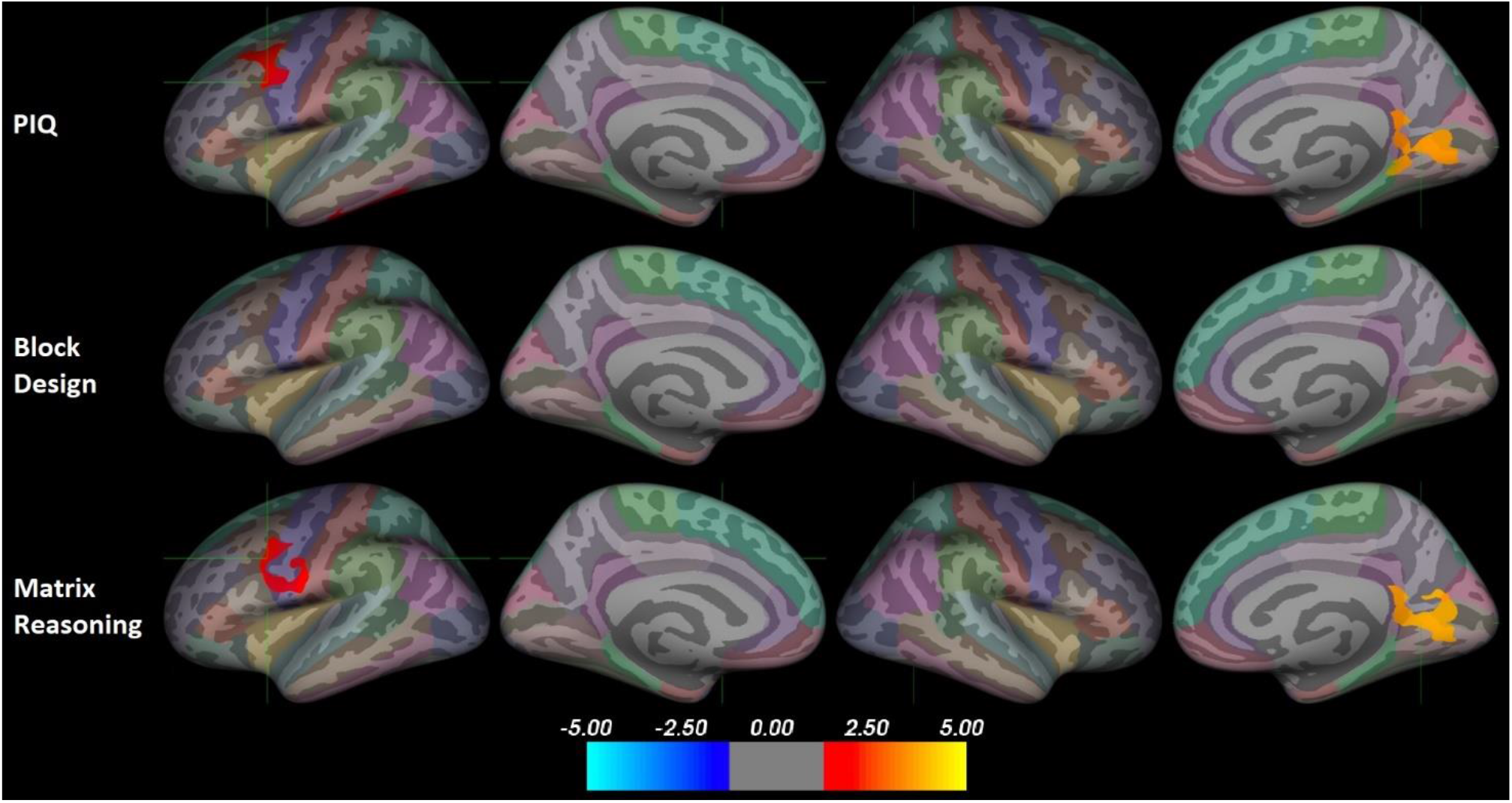
Positive associations between non-verbal ability and pial surface area. Results corrected for multiple comparisons using Monte-Carlo Null-Z simulation. Color indicates significance as a z-value. The position of the green crosshair indicates the most statistically significant vertex in statistically significant clusters. Left hemisphere on the left and right on the right side. Color coding of regions according to the Desikan–Killiany atlas. PIQ = performance intelligence quotient.

### Cortical thickness and non-verbal ability

Figure 3 presents the associations between CT and non-verbal ability. All significant associations were positive. There were no significant correlations between PIQ and CT. For Block Design only, there were significant clusters in the left precentral gyrus (peak z = 1.70, size = 959.0 mm^2^, peak coordinates -36.8, -18.3, 64.5) and the right postcentral gyrus (peak z = 2.26, size = 1158.2 mm^2^, peak coordinates 48.8, -16.5, 49.1). There were no significant correlations between Matrix Reasoning scores and CT.

**Figure 3.**
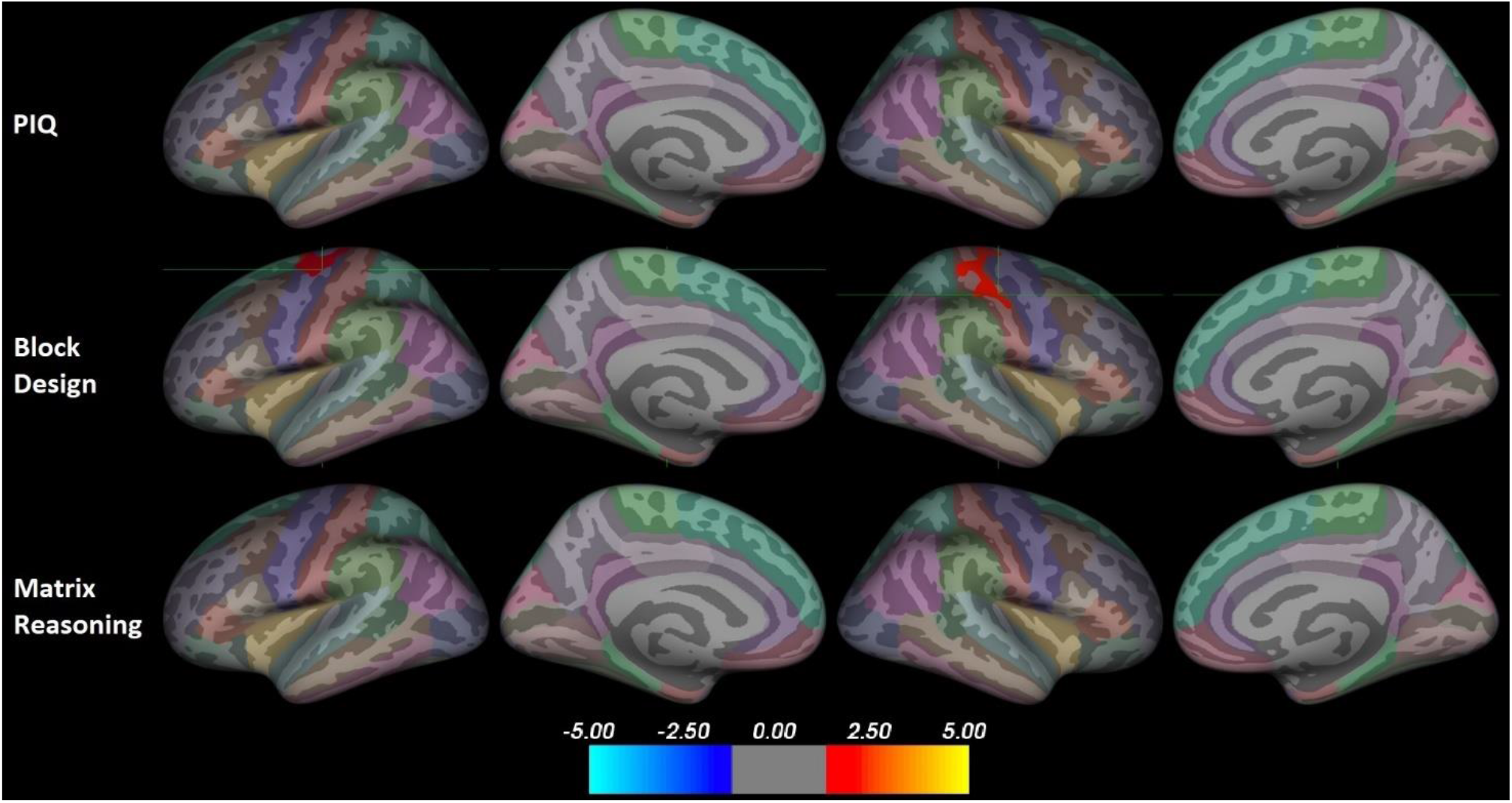
Positive associations between non-verbal ability and cortical thickness. Results corrected for multiple comparisons using Monte-Carlo Null-Z simulation. The position of the green crosshair indicates the most statistically significant vertex in statistically significant clusters. Left hemisphere on the left and right on the right side. Color coding of regions according to the Desikan–Killiany atlas. PIQ = performance intelligence quotient.

### Sensitivity analyses

The correlations between PIQ and brain metrics are shown in Supplementary Figures 2–7. Qdec images are presented without correction for multiple comparisons at z-value threshold 1.3. Overall, there are no large differences compared to the basic statistical model.

### ROI-based analyses

There were no correlations between non-verbal ability and either cortical volume, area, or thickness that survived correction for multiple comparisons. All correlations are presented in Supplementary Table.

### Post hoc analyses

We further visually explored the regions where PIQ associated with brain metrics to assess whether either subtest was driving the results. Supplementary Figure 8 presents the left caudal middle frontal gyrus associated with both volume and SA, as well as the left inferior temporal gyrus associated with SA. For the left caudal middle frontal gyrus, both subtests show some positive clusters in the same region. For the left inferior temporal gyrus, neither subtest alone shows large clusters in the region. In all three clusters, the results don’t seem to be strongly driven by either subtest.

Supplementary Figure 9 presents the right medial occipital region associated with both volume and SA. The Matrix Reasoning subtest shows large clusters in the same region for both volume and SA, while the Block Design subtest shows barely any small clusters in the region, suggesting that the correlations between PIQ and brain metrics in the right medial occipital region were driven by the Matrix Reasoning task performance.

### Discussion

In this study, we examined the associations between non-verbal ability and cortical brain structure (volume, SA, and CT) in a sample of typically developing 5-year-olds. We hypothesized based on the P-FIT model that non-verbal ability would be positively correlated with volume and SA in frontal and parietal regions. In line with the hypothesis, we found that the volume and SA of the left caudal middle frontal gyrus were positively associated with non-verbal ability. Additionally, we found significant positive associations with right medial occipital structure and left inferior temporal SA. Furthermore, we expected to find associations between non-verbal ability and CT in frontal and parietal regions. Two significant positive associations with visuospatial ability measures utilizing only one task were found but there were no associations with the overall non-verbal ability of the child. Altogether, this is the first study to examine the cortical structural correlates of non-verbal ability in a large sample of typically developing 5-year-olds. Our results suggest that some of the structures identified in studies of older participants are correlated with non-verbal ability at this stage of development.

We found associations between non-verbal ability and both volume and SA in the caudal middle frontal gyrus. The middle frontal gyrus is a region that is often associated with cognitive ability both structurally (Basten et al., 2015; Botdorf & Riggins, 2018; Brouwer et al., 2014; Frangou et al., 2004; Schilling et al., 2013) and functionally (Basten et al., 2015; Osaka et al., 2004), however the findings are typically seen in the frontal parts of the middle frontal gyrus. Furthermore, the pediatric studies on the topic have typically focused on CT rather than volume or SA (Botdorf & Riggins, 2018; Brouwer et al., 2014). The posterior parts of the caudal middle frontal gyrus form a part of the premotor cortex, which is, in addition to cognitive ability (Jung & Haier, 2007; O’Boyle et al., 2005), also a relevant region for mathematical ability (Navas-Sánchez et al., 2016), working memory (an fMRI study, Osaka et al., 2004), and speech perception (based on transcranial magnetic stimulation studies, see Meister et al., 2007; Sato et al., 2009). The premotor cortex is especially often observed relevant in functional brain studies focusing on cognitive ability (Jung & Haier, 2007; Osaka et al., 2004), while structural findings are comparatively scarce. Navas-Sánchez et al. (2016), observed larger SA in math gifted adolescents compared to high cognitive ability controls especially in the left caudal middle frontal gyrus. Some of the tests used in the study by Navas-Sánchez et al. (2016) were similar to ours, assessing visuospatial thinking and the ability to recognize patterns or rules, but they also measured other aspects of “mathematical giftedness”, such as intuition and creativity that may have completely different neurobiological correlates. Nevertheless, our results support the previous studies in proposing that the positive association between non-verbal ability and SA may already be observable at 5 years of age, which is in line with the finding that more intelligent children reach peak SA faster (Schnack et al., 2015).

Volume and SA in the right medial occipital region, including parts of the pericalcarine, isthmus of cingulate gyrus, precuneus, and lingual regions, were associated with non-verbal ability, and more specifically with visual abstract reasoning rather than visuospatial ability. Some studies in adults have found associations between general cognitive ability and the lingual gyrus volume (Colom et al., 2006b) as well as more widespread occipital GM volumes (Colom et al., 2006a; Haier et al., 2004). Notably, Colom et al. only found associations with visuospatial ability (visual abstract reasoning ability not tested, Colom et al., 2006a), while our results in the occipital lobe were driven by visual abstract reasoning ability. Furthermore, in previous articles, the associations between general cognitive ability and occipital brain metrics have generally been found on the lateral rather than medial surface in adults (Colom et al., 2006a) and children/adolescents (Karama et al., 2009). One study in children has found cortical thickening in left medial occipital cortex to be associated with higher visuospatial ability (Sowell et al., 2004). Parts of the occipital lobe are often involved in functional studies on cognitive function (Jung & Haier, 2007). For example, bilateral inferior occipital gyri activation is seen during deduction tasks (Goel & Dolan, 2004). Bilateral precuneus shows increased activation during non-verbal tasks in adolescents with high cognitive ability (Lee et al., 2006; O’Boyle et al., 2005). Notably, the activation is typically seen in more superior parts of the precuneus. To the best of our knowledge, this is the first study to link right medial occipital cortex volume and SA to non-verbal ability in children and thus, these areas should be included among the hypothesized structures related to non-verbal ability specifically.

In both volume and SA, we observed associations with visual abstract reasoning ability in largely similar areas than non-verbal ability. On the contrary, there were no associations between volume or SA and visuospatial ability. In contrast to our results, one previous study in adolescents measured both visual abstract reasoning and visuospatial ability (using the same subtests as we did) and found both to be associated with CT in the left frontal cortex, suggesting a common underlying neurobiology between visual abstract reasoning and visuospatial ability (Schilling et al., 2013). To further explore the possibility of a common neurobiological basis in our sample, we examined the Qdec analyses from the two subtests separately without correction for multiple comparisons. We observed a major difference between the clusters associated with visual abstract reasoning and visuospatial ability in the right medial occipital region, while the clusters in the prefrontal cortex were relatively similar. Our findings are not in conflict with previous findings that support the idea that the left middle frontal region is involved non-verbal ability (Navas-Sánchez et al., 2016; Schilling et al., 2013). On the other hand, our findings do suggest that the right medial occipital cortex volume and SA are associated with visual abstract reasoning ability but not with visuospatial ability. However, to the best of our knowledge, this is the first study to find this connection and further studies are needed to confirm the findings.

We found a positive association between non-verbal ability and SA in the inferior temporal gyrus. The inferior temporal gyrus, as well as the medial occipital region, has a key role in the ventral visual pathway (Kravitz et al., 2013), a network responsible for object recognition. On a related note, it has a role in the visual and auditory word processing (Cohen et al., 2004). In children with a family risk for dyslexia, a smaller SA was observed, even when controlling for the reading ability (Beelen et al., 2019). Additionally, one pediatric neuroimaging study found a positive association between general cognitive ability and inferior temporal CT in children (Karama et al., 2009). In theory, one would expect the structural and functional characteristics of the system responsible for object and pattern recognition to affect the performance on tasks that require pattern recognition (such as the tests in our study). However, current information on the roles of temporal and occipital cortices for non-verbal ability is conflicting (for review, please see Basten et al., 2015; Jung & Haier, 2007), and little is known about the role of these regions during childhood cognitive development.

We also found positive associations between CT and the visuospatial ability in the left precentral and right postcentral gyri. One previous study found widespread positive associations between Wechsler Abbreviated Scale of Intelligence (WASI) score and CT in 6-to 18-year-olds, also in older and younger halves separately (Karama et al., 2009). In their 6-to 12-year-old sample, the main overlap with our results is the positive association in the right postcentral gyrus. On the contrary, Botdorf & Riggins (2018) found no associations between general cognitive ability and CT in fronto–parietal regions but did find negative associations between CT and working memory (corrected for general cognitive ability) in multiple regions including the right postcentral gyrus in a sample of typically developing 4–8-olds. The primary somatosensory area is not commonly associated with the cognitive ability in structural neuroimaging studies but when it is, the findings tend to be on the right rather than on the left hemisphere (Haier et al., 2004; Jung & Haier, 2007; Karama et al., 2009). Decreasing CT in the left precentral gyrus has been associated with a decrease in general cognitive ability in a sample where children and adolescents were imaged twice approximately two years apart (Burgaleta et al., 2014), suggesting that too much thinning during childhood development may be associated with undesirable outcomes.

The most recent studies suggest the CT peaks at a very young age, possibly even before 2 years of age (Bethlehem et al., 2022; Frangou et al., 2022). Therefore, in a simplistic “more advanced is better” interpretation, the participants with higher non-verbal ability would be further in the developmental trajectory and have lower CT. Some studies have indeed found higher general cognitive ability (Schnack et al., 2015; Squeglia et al., 2013) and working memory (Botdorf & Riggins, 2018) to be associated with lower CT in children and adolescents. However, the positive associations seen in our study, while in agreement with many previous studies (Girault et al., 2020; Leonard et al., 2019; Meruelo et al., 2019; Schilling et al., 2013), contrast the idea that more advanced development would necessarily correlate with higher cognitive ability. One option to consider is that individuals may have different growth trajectories depending on cognitive ability. Shaw et al. (2006) have shown that the children with higher general cognitive ability reach their peak CT later, while Khundrakpam et al. (2017) suggest different CT coupling between the cortical regions between ages 6–18 years based on verbal ability. Meanwhile, other studies have found positive associations between general cognitive ability and CT in multiple brain regions in 6–18-year-olds (Karama et al., 2009, 2011), 9–24-year-olds (Menary et al., 2013), and adults (Bajaj et al., 2018) suggesting that individuals with higher general cognitive ability may retain greater CT, although there have been conflicting results in adult studies, too (Tadayon et al., 2020). Overall, the results regarding cognitive ability and CT in children are currently inconsistent and more studies are needed. There are currently some large multisite neuroimaging projects devoted to longitudinal data collection of the developing brain, such as the HEALthy Brain and Child Development consortium (HBCD; Volkow et al., 2021) and the Adolescent Brain Cognitive Development consortium (ABCD; Hagler et al., 2019; Volkow et al., 2018), which will provide crucial information on developmental trajectories of the brain.

Both a strength and a limitation of this study is the limited age range in a cross-sectional setting. While the strength lies in the possibility to understand the neural correlates of non-verbal ability at this specific age relevant for later development, it precludes true longitudinal and developmental interpretations. Especially with CT, it seems to be the case that longitudinal modeling is needed to find the potential individual differences in growth trajectories and how they might relate to non-verbal ability. On the other hand, to our knowledge, this is the first neuroimaging study to explore association between non-verbal ability and structural brain development in a large sample of typically developing 5-year-olds. The small age range provides an opportunity to get an accurate image of brain structure at this stage of development. Another limitation is the generalizability of the results especially to clinical samples. The participants in the final sample were born at a higher gestational age and had less visits to the NICU, suggesting that many participants with even slight issues during pregnancy or perinatal period were not included in the sample. Furthermore, our sample is highly ethnically homogenous, and the results are not necessarily generalizable to non-Caucasian populations.

## Conclusion

To the best of our knowledge, this is the first study to explore cortical structural development in relation to non-verbal ability in a large sample of typically developing 5-year-olds. We found that non-verbal ability was associated with volume and SA in left middle frontal and right medial occipital regions, and especially the medial occipital region was associated with visual abstract reasoning rather than visuospatial ability. On the other hand, CT in left precentral and right postcentral gyri were only associated with visuospatial ability specifically. Discrepancy between CT results and other results is not surprising considering that CT develops relatively independently from SA and volume (Winkler et al., 2010) and has a different genetic basis (Panizzon et al., 2009). Most associations between brain structure and non-verbal ability were found in frontoparietal regions as expected based on the P-FIT model (Jung & Haier, 2007), the most notable exception being the right medial occipital region. All associations were positive, which was also in line with previous literature in pediatric populations. Overall, neural characteristics of cognitive development should be studied in samples of different ages and backgrounds. Longitudinal studies involving young children will be especially important to characterize the potential individual differences in developmental trajectories.

## Acknowledgements

This work was supported by Päivikki and Sakari Sohlberg Foundation (to E.P.P.), Juho Vainio Foundation (to E.P.P.), Emil Aaltonen Foundation (to E.P.P, to A.C.), Turku University Foundation (to A.C.), the Finnish Medical Foundation (#5303 to V.K.), the Finnish Cultural Foundation (#00190572 to V.K.), Finnish Brain Foundation (to E.Si.), the Academy of Finland (#26080983 to H.M., #325292 to L.K.), Signe and Ane Gyllenberg Foundation (to L.K., H.K.), and Finnish State Grants for Clinical Research (ERVA P3654 to L.K., to H.K.).

We thank our research nurse Susanne Sinisalo for her expertise in study management and performing the scans with the investigators and all participating FinnBrain Families.

The authors declare no conflicts of interest.

## References

Aoki, H., Fujino, M., Arai, I., Yasuhara, H., Ebisu, R., Ohgitani, A., & Minowa, H. (2020). The efficacy of routine brain MRI for term neonates admitted to neonatal intensive care unit. The Journal of Maternal-Fetal & Neonatal Medicine, 35(15), 2932–2935. https://doi.org/10.1080/14767058.2020.1814240

Aubert-Broche, B., Fonov, V. S., García-Lorenzo, D., Mouiha, A., Guizard, N., Coupé, P., Eskildsen, S. F., & Collins, D. L. (2013). A new method for structural volume analysis of longitudinal brain MRI data and its application in studying the growth trajectories of anatomical brain structures in childhood. NeuroImage, 82, 393–402. https://doi.org/10.1016/J.NEUROIMAGE.2013.05.065

Bajaj, S., Raikes, A., Smith, R., Dailey, N. S., Alkozei, A., Vanuk, J. R., & Killgore, W. D. S. (2018). The Relationship Between General Intelligence and Cortical Structure in Healthy Individuals. Neuroscience, 388, 36–44. https://doi.org/10.1016/J.NEUROSCIENCE.2018.07.008

Basten, U., Hilger, K., & Fiebach, C. J. (2015). Where smart brains are different: A quantitative meta-analysis of functional and structural brain imaging studies on intelligence. Intelligence, 51, 10–27. https://doi.org/10.1016/J.INTELL.2015.04.009

Batty, G. D., Deary, I. J., & Gottfredson, L. S. (2007). Premorbid (early life) IQ and Later Mortality Risk: Systematic Review. Annals of Epidemiology, 17(4), 278–288. https://doi.org/10.1016/J.ANNEPIDEM.2006.07.010

Batty, G. D., Deary, I. J., & Macintyre, S. (2006). Childhood IQ and life course socioeconomic position in relation to alcohol induced hangovers in adulthood: the Aberdeen children of the 1950s study. Journal of Epidemiology & Community Health, 60(10), 872–874. https://doi.org/10.1136/JECH.2005.045039

Beelen, C., Vanderauwera, J., Wouters, J., Vandermosten, M., & Ghesquière, P. (2019). Atypical gray matter in children with dyslexia before the onset of reading instruction. Cortex, 121, 399–413. https://doi.org/10.1016/J.CORTEX.2019.09.010

Bertua, C., Anderson, N., & Salgado, J. F. (2005). The predictive validity of cognitive ability tests: A UK meta-analysis. Journal of Occupational and Organizational Psychology, 78(3), 387–409. https://doi.org/10.1348/096317905X26994

Bethlehem, R. A. I., Seidlitz, J., White, S. R., Vogel, J. W., Anderson, K. M., Adamson, C., Adler, S., Alexopoulos, G. S., Anagnostou, E., Areces-Gonzalez, A., Astle, D. E., Auyeung, B., Ayub, M., Bae, J., Ball, G., Baron-Cohen, S., Beare, R., Bedford, S. A., Benegal, V.,… Alexander-Bloch, A. F. (2022). Brain charts for the human lifespan. Nature 2022 604:7906, 604(7906), 525–533. https://doi.org/10.1038/S41586-022-04554-Y

Botdorf, M., & Riggins, T. (2018). When less is more: Thinner fronto-parietal cortices are associated with better forward digit span performance during early childhood. Neuropsychologia, 121, 11–18. https://doi.org/10.1016/J.NEUROPSYCHOLOGIA.2018.10.020

Brouwer, R. M., Soelen, I. L. C. van, Swagerman, S. C., Schnack, H. G., Ehli, E. A., Kahn, R. S., Pol, H. E. H., & Boomsma, D. I. (2014). Genetic associations between intelligence and cortical thickness emerge at the start of puberty. Human Brain Mapping, 35(8), 3760–3773. https://doi.org/10.1002/HBM.22435

Brown, M. I., Wai, J., & Chabris, C. F. (2021). Can You Ever Be Too Smart for Your Own Good? Comparing Linear and Nonlinear Effects of Cognitive Ability on Life Outcomes. Perspectives on Psychological Science, 16(6), 1337–1359. https://doi.org/10.1177/1745691620964122

Brown, T. T., Kuperman, J. M., Chung, Y., Erhart, M., McCabe, C., Hagler, D. J., Venkatraman, V. K., Akshoomoff, N., Amaral, D. G., Bloss, C. S., Casey, B. J., Chang, L., Ernst, T. M., Frazier, J. A., Gruen, J. R., Kaufmann, W. E., Kenet, T., Kennedy, D. N., Murray, S. S., … Dale, A. M. (2012). Neuroanatomical Assessment of Biological Maturity. Current Biology, 22(18), 1693–1698. https://doi.org/10.1016/J.CUB.2012.07.002

Burgaleta, M., Johnson, W., Waber, D. P., Colom, R., & Karama, S. (2014). Cognitive ability changes and dynamics of cortical thickness development in healthy children and adolescents. NeuroImage, 84, 810–819. https://doi.org/10.1016/J.NEUROIMAGE.2013.09.038

Chandola, T., Deary, I. J., Blane, D., & Batty, G. D. (2006). Childhood IQ in relation to obesity and weight gain in adult life: The National Child Development (1958) Study. International Journal of Obesity, 30(9), 1422–1432. https://doi.org/10.1038/sj.ijo.0803279

Chang, L., Oishi, K., Skranes, J., Buchthal, S., Cunningham, E., Yamakawa, R., Hayama, S., Jiang, C. S., Alicata, D., Hernandez, A., Cloak, C., Wright, T., & Ernst, T. (2016). Sex-Specific Alterations of White Matter Developmental Trajectories in Infants With Prenatal Exposure to Methamphetamine and Tobacco. JAMA Psychiatry, 73(12), 1217. https://doi.org/10.1001/jamapsychiatry.2016.2794

Chyl, K., Fraga-González, G., Brem, S., & Jednoróg, K. (2021). Brain dynamics of (a)typical reading development—a review of longitudinal studies. NPJ Science of Learning, 6(1). https://doi.org/10.1038/S41539-020-00081-5

Cohen, L., Jobert, A., Le Bihan, D., & Dehaene, S. (2004). Distinct unimodal and multimodal regions for word processing in the left temporal cortex. NeuroImage, 23(4), 1256–1270. https://doi.org/10.1016/J.NEUROIMAGE.2004.07.052

Colom, R., Jung, R. E., & Haier, R. J. (2006a). Distributed brain sites for the g-factor of intelligence. NeuroImage, 31(3), 1359–1365. https://doi.org/10.1016/J.NEUROIMAGE.2006.01.006

Colom, R., Jung, R. E., & Haier, R. J. (2006b). Finding the g-factor in brain structure using the method of correlated vectors. Intelligence, 34(6), 561–570. https://doi.org/10.1016/J.INTELL.2006.03.006

Copeland, A. M., Silver, E. A. O., Korja, R., Lehtola, S., Merisaari, H., Saukko, E., Sinisalo, S., Saunavaara, J., Lähdesmäki, T., Parkkola, R., Nolvi, S., Karlsson, L., Karlsson, H., & Tuulari, J. (2021). Infant and child MRI: a review of scanning procedures. Frontiers in Neuroscience, 15, 632. https://doi.org/10.3389/FNINS.2021.666020

Courchesne, E., Chisum, H. J., Townsend, J., Cowles, A., Covington, J., Egaas, B., Harwood, M., Hinds, S., & Press, G. A. (2000). Normal Brain Development and Aging: Quantitative Analysis at in Vivo MR Imaging in Healthy Volunteers1. Https://Doi-Org.Ezproxy.Utu.Fi/10.1148/Radiology.216.3.R00au37672, 216(3), 672–682. https://doi.org/10.1148/RADIOLOGY.216.3.R00AU37672

Deary, I. J., Pattie, A., & Starr, J. M. (2013). The Stability of Intelligence From Age 11 to Age 90 Years: The Lothian Birth Cohort of 1921. Psychological Science, 24(12), 2361–2368. https://doi.org/10.1177/0956797613486487/ASSET/IMAGES/LARGE/10.1177_0956797613486487-FIG2.JPEG

Deary, I. J., Spinath, F. M., & Bates, T. C. (2006). Genetics of intelligence. In European Journal of Human Genetics (Vol. 14, Issue 6, pp. 690–700). Nature Publishing Group. https://doi.org/10.1038/sj.ejhg.5201588

Deary, I. J., Strand, S., Smith, P., & Fernandes, C. (2007). Intelligence and educational achievement. Intelligence, 35(1), 13–21. https://doi.org/10.1016/J.INTELL.2006.02.001

Desikan, R. S., Ségonne, F., Fischl, B., Quinn, B. T., Dickerson, B. C., Blacker, D., Buckner, R. L., Dale, A. M., Maguire, R. P., Hyman, B. T., Albert, M. S., & Killiany, R. J. (2006). An automated labeling system for subdividing the human cerebral cortex on MRI scans into gyral based regions of interest. NeuroImage, 31(3), 968–980. https://doi.org/10.1016/j.neuroimage.2006.01.021

Donald, K. A., Eastman, E., Howells, F. M., Adnams, C., Riley, E. P., Woods, R. P., Narr, K. L., & Stein, D. J. (2015). Neuroimaging effects of prenatal alcohol exposure on the developing human brain: a magnetic resonance imaging review. Acta Neuropsychiatrica, 27(05), 251–269. https://doi.org/10.1017/neu.2015.12

Faridi, N., Karama, S., Burgaleta, M., White, M. T., Evans, A. C., Fonov, V., Collins, D. L., & Waber, D. P. (2015). Neuroanatomical correlates of behavioral rating versus performance measures of working memory in typically developing children and adolescents. Neuropsychology, 29(1), 82–91. https://doi.org/10.1037/neu0000079

Fischl, B., & Dale, A. M. (2000). Measuring the thickness of the human cerebral cortex from magnetic resonance images. Proceedings of the National Academy of Sciences of the United States of America, 97(20), 11050–11055. https://doi.org/10.1073/pnas.200033797

Fischl, B., Sereno, M. I., & Dale, A. M. (1999). Cortical surface-based analysis: II. Inflation, flattening, and a surface-based coordinate system. NeuroImage, 9(2), 195–207. https://doi.org/10.1006/nimg.1998.0396

Fischl, B., Sereno, M. I., Tootell, R. B. H., & Dale, A. M. (1999). High-resolution intersubject averaging and a coordinate system for the cortical surface. Human Brain Mapping, 8(4), 272–284. https://doi.org/10.1002/(SICI)1097-0193(1999)8:4<272::AID-HBM10>3.0.CO;2-4

Frangou, S., Chitins, X., & Williams, S. C. R. (2004). Mapping IQ and gray matter density in healthy young people. NeuroImage, 23(3), 800–805. https://doi.org/10.1016/J.NEUROIMAGE.2004.05.027

Frangou, S., Modabbernia, A., Williams, S. C. R., Papachristou, E., Doucet, G. E., Agartz, I., Aghajani, M., Akudjedu, T. N., Albajes-Eizagirre, A., Alnæs, D., Alpert, K. I., Andersson, M., Andreasen, N. C., Andreassen, O. A., Asherson, P., Banaschewski, T., Bargallo, N., Baumeister, S., Baur-Streubel, R., … Dima, D. (2022). Cortical thickness across the lifespan: Data from 17,075 healthy individuals aged 3– 90 years. Human Brain Mapping, 43(1), 431–451. https://doi.org/10.1002/HBM.25364

Furnham, A., & Cheng, H. (2017). Childhood Cognitive Ability Predicts Adult Financial Well-Being. Journal of Intelligence, 5(1), 1–12. https://doi.org/10.3390/JINTELLIGENCE5010003

Girault, J. B., Cornea, E., Goldman, B. D., Jha, S. C., Murphy, V. A., Li, G., Wang, L., Shen, D., Knickmeyer, R. C., Styner, M., & Gilmore, J. H. (2020). Cortical Structure and Cognition in Infants and Toddlers. Cerebral Cortex, 30(2), 786–800. https://doi.org/10.1093/CERCOR/BHZ126

Goel, V., & Dolan, R. J. (2004). Differential involvement of left prefrontal cortexin inductive and deductive reasoning. Cognition, 93(3), B109–B121. https://doi.org/10.1016/J.COGNITION.2004.03.001

Gow, A. J., Johnson, W., Pattie, A., Brett, C. E., Roberts, B., Starr, J. M., & Deary, I. J. (2011). Stability and Change in Intelligence From Age 11 to Ages 70, 79, and 87: The Lothian Birth Cohorts of 1921 and 1936. Psychology and Aging, 26(1), 232–240. https://doi.org/10.1037/a0021072

Hagler, D. J., Hatton, S. N., Cornejo, M. D., Makowski, C., Fair, D. A., Dick, A. S., Sutherland, M. T., Casey, B. J., Barch, D. M., Harms, M. P., Watts, R., Bjork, J. M., Garavan, H. P., Hilmer, L., Pung, C. J., Sicat, C. S., Kuperman, J., Bartsch, H., Xue, F., … Dale, A. M. (2019). Image processing and analysis methods for the Adolescent Brain Cognitive Development Study. NeuroImage, 202, 116091. https://doi.org/10.1016/J.NEUROIMAGE.2019.116091

Hagler, D. J., Saygin, A. P., & Sereno, M. I. (2006). Smoothing and cluster thresholding for cortical surface-based group analysis of fMRI data. NeuroImage, 33(4), 1093–1103. https://doi.org/10.1016/J.NEUROIMAGE.2006.07.036

Haier, R. J., Jung, R. E., Yeo, R. A., Head, K., & Alkire, M. T. (2004). Structural brain variation and general intelligence. NeuroImage, 23(1), 425–433. https://doi.org/10.1016/J.NEUROIMAGE.2004.04.025

Hashimoto, T., Matsuzaki, Y., Yokota, S., & Kawashima, R. (2022). Academic achievements and brain volume development in children and adolescents. Cerebral Cortex Communications, 3(4). https://doi.org/10.1093/TEXCOM/TGAC048

Haworth, C. M. A., Wright, M. J., Luciano, M., Martin, N. G., de Geus, E. J. C., van Beijsterveldt, C. E. M., Bartels, M., Posthuma, D., Boomsma, D. I., Davis, O. S. P., Kovas, Y., Corley, R. P., DeFries, J. C., Hewitt, J. K., Olson, R. K., Rhea, S.-A., Wadsworth, S. J., Iacono, W. G., McGue, M., … Plomin, R. (2009). The heritability of general cognitive ability increases linearly from childhood to young adulthood. Molecular Psychiatry 2010 15:11, 15(11), 1112–1120. https://doi.org/10.1038/MP.2009.55

Hong, H. S., & Lee, J. Y. (2018). Intracranial hemorrhage in term neonates. Child’s Nervous System, 34(6), 1135–1143. https://doi.org/10.1007/S00381-018-3788-8/TABLES/4

JASP Team. (2022). JASP (Version 0.16.1)[Computer software]. https://jasp-stats.org/

Jeong, H. J., Shim, S.-Y., Cho, H. J., Cho, S. J., Son, D. W., & Park, E. A. (2016). Cerebellar Development in Preterm Infants at Term-Equivalent Age Is Impaired after Low-Grade Intraventricular Hemorrhage. The Journal of Pediatrics. https://doi.org/10.1016/j.jpeds.2016.05.010

Jha, S. C., Xia, K., Schmitt, J. E., Ahn, M., Girault, J. B., Murphy, V. A., Li, G., Wang, L., Shen, D., Zou, F., Zhu, H., Styner, M., Knickmeyer, R. C., & Gilmore, J. H. (2018). Genetic influences on neonatal cortical thickness and surface area. Human Brain Mapping, 39(12), 4998–5013. https://doi.org/10.1002/HBM.24340

Jung, R. E., & Haier, R. J. (2007). The Parieto-Frontal Integration Theory (P-FIT) of intelligence: Converging neuroimaging evidence. Behavioral and Brain Sciences, 30(2), 135–154. https://doi.org/10.1017/S0140525X07001185

Kapellou, O., Counsell, S. J., Kennea, N., Dyet, L., Saeed, N., Stark, J., Maalouf, E., Duggan, P., Ajayi-Obe, M., Hajnal, J., Allsop, J. M., Boardman, J., Rutherford, M. A., Cowan, F., & Edwards, A. D. (2006). Abnormal Cortical Development after Premature Birth Shown by Altered Allometric Scaling of Brain Growth. PLOS Medicine, 3(8), e265. https://doi.org/10.1371/journal.pmed.0030265

Karama, S., Ad-Dab’bagh, Y., Haier, R. J., Deary, I. J., Lyttelton, O. C., Lepage, C., & Evans, A. C. (2009). Positive association between cognitive ability and cortical thickness in a representative US sample of healthy 6 to 18 year-olds. Intelligence, 37(2), 145–155. https://doi.org/10.1016/J.INTELL.2008.09.006

Karama, S., Colom, R., Johnson, W., Deary, I. J., Haier, R., Waber, D. P., Lepage, C., Ganjavi, H., Jung, R., & Evans, A. C. (2011). Cortical thickness correlates of specific cognitive performance accounted for by the general factor of intelligence in healthy children aged 6 to 18. NeuroImage, 55(4), 1443–1453. https://doi.org/10.1016/J.NEUROIMAGE.2011.01.016

Karlsson, L., Tolvanen, M., Scheinin, N. M., Uusitupa, H.-M., Korja, R., Ekholm, E., Tuulari, J. J., Pajulo, M., Huotilainen, M., Paunio, T., & Karlsson, H. (2018). Cohort Profile: The FinnBrain Birth Cohort Study (FinnBrain). International Journal of Epidemiology, 47(1), 15–16j. https://doi.org/10.1093/ije/dyx173

Khundrakpam, B. S., Lewis, J. D., Reid, A., Karama, S., Zhao, L., Chouinard-Decorte, F., & Evans, A. C. (2017). Imaging structural covariance in the development of intelligence. NeuroImage, 144, 227–240. https://doi.org/10.1016/J.NEUROIMAGE.2016.08.041

Kim, D. J., Davis, E. P., Sandman, C. A., Sporns, O., O’Donnell, B. F., Buss, C., & Hetrick, W. P. (2016). Children’s intellectual ability is associated with structural network integrity. NeuroImage, 124, 550–556. https://doi.org/10.1016/J.NEUROIMAGE.2015.09.012

Knickmeyer, R. C., Gouttard, S., Kang, C., Evans, D., Wilber, K., Smith, J. K., Hamer, R. M., Lin, W., Gerig, G., & Gilmore, J. H. (2008). A structural MRI study of human brain development from birth to 2 years. The Journal of Neuroscience : The Official Journal of the Society for Neuroscience, 28(47), 12176–12182. https://doi.org/10.1523/jneurosci.3479-08.2008

Knickmeyer, R. C., Xia, K., Lu, Z., Ahn, M., Jha, S. C., Zou, F., Zhu, H., Styner, M., & Gilmore, J. H. (2016). Impact of Demographic and Obstetric Factors on Infant Brain Volumes: A Population Neuroscience Study. Cerebral Cortex. https://doi.org/10.1093/cercor/bhw331

Kravitz, D. J., Saleem, K. S., Baker, C. I., Ungerleider, L. G., & Mishkin, M. (2013). The ventral visual pathway: An expanded neural framework for the processing of object quality. Trends in Cognitive Sciences, 17(1), 26. https://doi.org/10.1016/J.TICS.2012.10.011

Kumpulainen, V., Lehtola, S. J., Tuulari, J. J., Silver, E., Copeland, A., Korja, R., Karlsson, H., Karlsson, L., Merisaari, H., Parkkola, R., Saunavaara, J., Lähdesmäki, T., & Scheinin, N. M. (2020). Prevalence and Risk Factors of Incidental Findings in Brain MRIs of Healthy Neonates—The FinnBrain Birth Cohort Study. Frontiers in Neurology, 10, 1347. https://doi.org/10.3389/fneur.2019.01347

Kuperberg, G. R., Broome, M. R., McGuire, P. K., David, A. S., Eddy, M., Ozawa, F., Goff, D., West, W. C., Williams, S. C. R., Van der Kouwe, A. J. W., Salat, D. H., Dale, A. M., & Fischl, B. (2003). Regionally localized thinning of the cerebral cortex in schizophrenia. Archives of General Psychiatry, 60(9), 878–888. https://doi.org/10.1001/archpsyc.60.9.878

Lang, J. W. B., & Kell, H. J. (2020). General mental ability and specific abilities: Their relative importance for extrinsic career success. Journal of Applied Psychology, 105(9), 1047–1061. https://doi.org/10.1037/apl0000472

Lee, K. H., Choi, Y. Y., Gray, J. R., Cho, S. H., Chae, J. H., Lee, S., & Kim, K. (2006). Neural correlates of superior intelligence: Stronger recruitment of posterior parietal cortex. NeuroImage, 29(2), 578–586. https://doi.org/10.1016/J.NEUROIMAGE.2005.07.036

Lenroot, R. K., Schmitt, J. E., Ordaz, S. J., Wallace, G. L., Neale, M. C., Lerch, J. P., Kendler, K. S., Evans, A. C., & Giedd, J. N. (2009). Differences in genetic and environmental influences on the human cerebral cortex associated with development during childhood and adolescence. Human Brain Mapping, 30(1), 163–174. https://doi.org/10.1002/HBM.20494

Leonard, J. A., Romeo, R. R., Park, A. T., Takada, M. E., Robinson, S. T., Grotzinger, H., Last, B. S., Finn, A. S., Gabrieli, J. D. E., & Mackey, A. P. (2019). Associations between cortical thickness and reasoning differ by socioeconomic status in development. Developmental Cognitive Neuroscience, 36, 100641. https://doi.org/10.1016/j.dcn.2019.100641

Li, X., Andres, A., Shankar, K., Pivik, R. T., Glasier, C. M., Ramakrishnaiah, R. H., Zhang, Y., Badger, T. M., & Ou, X. (2016). Differences in brain functional connectivity at resting state in neonates born to healthy obese or normal-weight mothers. Int J Obes, 40(12), 1931–1934. https://doi.org/10.1038/ijo.2016.166

Meister, I. G., Wilson, S. M., Deblieck, C., Wu, A. D., & Iacoboni, M. (2007). The Essential Role of Premotor Cortex in Speech Perception. Current Biology, 17(19), 1692–1696. https://doi.org/10.1016/J.CUB.2007.08.064

Menary, K., Collins, P. F., Porter, J. N., Muetzel, R., Olson, E. A., Kumar, V., Steinbach, M., Lim, K. O., & Luciana, M. (2013). Associations between cortical thickness and general intelligence in children, adolescents and young adults. Intelligence, 41(5), 597. https://doi.org/10.1016/J.INTELL.2013.07.010

Merisaari, H., Tuulari, J. J., Karlsson, L., Scheinin, N. M., Parkkola, R., Saunavaara, J., Lähdesmäki, T., Lehtola, S. J., Keskinen, M., Lewis, J. D., Evans, A. C., & Karlsson, H. (2019). Test-retest reliability of Diffusion Tensor Imaging metrics in neonates. NeuroImage, 197, 598–607. https://doi.org/10.1016/j.neuroimage.2019.04.067

Meruelo, A. D., Jacobus, J., Idy, E., Nguyen-Louie, T., Brown, G., & Tapert, S. F. (2019). Early adolescent brain markers of late adolescent academic functioning. Brain Imaging and Behavior, 13(4), 945–952. https://doi.org/10.1007/S11682-018-9912-2

Navas-Sánchez, F. J., Carmona, S., Alemán-Gómez, Y., Sánchez-González, J., Guzmán-de-Villoria, J., Franco, C., Robles, O., Arango, C., & Desco, M. (2016). Cortical morphometry in frontoparietal and default mode networks in math-gifted adolescents. Human Brain Mapping, 37(5), 1893–1902. https://doi.org/10.1002/hbm.23143

Neisser, U., Boodoo, G., Bouchard, T. J., Boykin, A. W., Brody, N., Ceci, S. J., Halpern, D. F., Loehlin, J. C., Perloff, R., Sternberg, R. J., & Urbina, S. (1996). Intelligence: Knowns and Unknowns. American Psychologist, 51(2), 77–101. https://doi.org/10.1037/0003-066X.51.2.77

O’Boyle, M. W., Cunnington, R., Silk, T. J., Vaughan, D., Jackson, G., Syngeniotis, A., & Egan, G. F. (2005). Mathematically gifted male adolescents activate a unique brain network during mental rotation. Cognitive Brain Research, 25(2), 583–587. https://doi.org/10.1016/J.COGBRAINRES.2005.08.004

Osaka, N., Osaka, M., Kondo, H., Morishita, M., Fukuyama, H., & Shibasaki, H. (2004). The neural basis of executive function in working memory: an fMRI study based on individual differences. NeuroImage, 21(2), 623–631. https://doi.org/10.1016/J.NEUROIMAGE.2003.09.069

Ou, X., Thakali, K. M., Shankar, K., Andres, A., & Badger, T. M. (2015). Maternal adiposity negatively influences infant brain white matter development: Maternal Obesity and Infant Brain. Obesity, 23(5), 1047–1054. https://doi.org/10.1002/oby.21055

Pangelinan, M. M., Zhang, G., VanMeter, J. W., Clark, J. E., Hatfield, B. D., & Haufler, A. J. (2011). Beyond age and gender: Relationships between cortical and subcortical brain volume and cognitive-motor abilities in school-age children. NeuroImage, 54(4), 3093–3100. https://doi.org/10.1016/J.NEUROIMAGE.2010.11.021

Panizzon, M. S., Fennema-Notestine, C., Eyler, L. T., Jernigan, T. L., Prom-Wormley, E., Neale, M., Jacobson, K., Lyons, M. J., Grant, M. D., Franz, C. E., Xian, H., Tsuang, M., Fischl, B., Seidman, L., Dale, A., & Kremen, W. S. (2009). Distinct Genetic Influences on Cortical Surface Area and Cortical Thickness. Cerebral Cortex, 19(11), 2728–2735. https://doi.org/10.1093/CERCOR/BHP026

Phan, T. V., Smeets, D., Talcott, J. B., & Vandermosten, M. (2018). Processing of structural neuroimaging data in young children: Bridging the gap between current practice and state-of-the-art methods. In Developmental Cognitive Neuroscience (Vol. 33, pp. 206–223). Elsevier Ltd. https://doi.org/10.1016/j.dcn.2017.08.009

Plomin, R., Fulker, D. W., Corley, R., & DeFries, J. C. (1997). Nature, Nurture, and Cognitive Development from 1 to 16 Years: A Parent-Offspring Adoption Study. Psychological Science, 8(6), 442–447. https://doi.org/10.1111/J.1467-9280.1997.TB00458.X

Plomin, R., & Von Stumm, S. (2018). The new genetics of intelligence. In Nature Reviews Genetics (Vol. 19, Issue 3, pp. 148–159). Nature Publishing Group. https://doi.org/10.1038/nrg.2017.104

Posthuma, D., De Geus, E. J. C., Baaré, W. F. C., Hulshoff Pol, H. E., Kahn, R. S., & Boomsma, D. I. (2002). The association between brain volume and intelligence is of genetic origin. In Nature Neuroscience (Vol. 5, Issue 2, pp. 83–84). Nature Publishing Group. https://doi.org/10.1038/nn0202-83

Pulli, E. P., Silver, E., Kumpulainen, V., Copeland, A., Merisaari, H., Saunavaara, J., Parkkola, R., Lähdesmäki, T., Saukko, E., Nolvi, S., Kataja, E.-L., Korja, R., Karlsson, L., Karlsson, H., & Tuulari, J. J. (2022). Feasibility of FreeSurfer Processing for T1-Weighted Brain Images of 5-Year-Olds: Semiautomated Protocol of FinnBrain Neuroimaging Lab. Frontiers in Neuroscience, 0, 640. https://doi.org/10.3389/FNINS.2022.874062

Qi, T., Schaadt, G., & Friederici, A. D. (2019). Cortical thickness lateralization and its relation to language abilities in children. Developmental Cognitive Neuroscience, 39, 100704. https://doi.org/10.1016/j.dcn.2019.100704

Ramsden, S., Richardson, F. M., Josse, G., Thomas, M. S. C., Ellis, C., Shakeshaft, C., Seghier, M. L., & Price, C. J. (2011). Verbal and nonverbal intelligence changes in the teenage brain. Nature, 479(7371), 113. https://doi.org/10.1038/NATURE10514

Raznahan, A., Shaw, P., Lalonde, F., Stockman, M., Wallace, G. L., Greenstein, D., Clasen, L., Gogtay, N., & Giedd, J. N. (2011). How Does Your Cortex Grow? Journal of Neuroscience, 31(19), 7174–7177. https://doi.org/10.1523/JNEUROSCI.0054-11.2011

Reiss, A. L., Abrams, M. T., Singer, H. S., Ross, J. L., & Denckla, M. B. (1996). Brain development, gender and IQ in children: A volumetric imaging study. Brain, 119(5), 1763–1774. https://doi.org/10.1093/BRAIN/119.5.1763

Rosas, H. D., Liu, A. K., Hersch, S., Glessner, M., Ferrante, R. J., Salat, D. H., Van Der Kouwe, A., Jenkins, B. G., Dale, A. M., & Fischl, B. (2002). Regional and progressive thinning of the cortical ribbon in Huntington’s disease. Neurology, 58(5), 695–701. https://doi.org/10.1212/WNL.58.5.695

Rosberg, A., Tuulari, J. J., Kumpulainen, V., Lukkarinen, M., Pulli, E. P., Silver, E., Copeland, A., Saukko, E., Saunavaara, J., Lewis, J. D., Karlsson, L., Karlsson, H., & Merisaari, H. (2022). Test–retest reliability of diffusion tensor imaging scalars in 5-year-olds. Human Brain Mapping, 43(16), 4984–4994. https://doi.org/10.1002/HBM.26064

Salat, D. H. (2004). Thinning of the Cerebral Cortex in Aging. Cerebral Cortex, 14(7), 721–730. https://doi.org/10.1093/cercor/bhh032

Sato, M., Tremblay, P., & Gracco, V. L. (2009). A mediating role of the premotor cortex in phoneme segmentation. Brain and Language, 111(1), 1–7. https://doi.org/10.1016/J.BANDL.2009.03.002

Schilling, C., Kühn, S., Paus, T., Romanowski, A., Banaschewski, T., Barbot, A., Barker, G. J., Brühl, R., Büchel, C., Conrod, P. J., Dalley, J. W., Flor, H., Ittermann, B., Ivanov, N., Mann, K., Martinot, J. L., Nees, F., Rietschel, M., Robbins, T. W., … Gallinat, J. (2013). Cortical thickness of superior frontal cortex predicts impulsiveness and perceptual reasoning in adolescence. Molecular Psychiatry, 18(5), 624–630. https://doi.org/10.1038/mp.2012.56

Schmidt, F. L., & Hunter, J. (2004). General Mental Ability in the World of Work: Occupational Attainment and Job Performance. Journal of Personality and Social Psychology, 86(1), 162–173. https://doi.org/10.1037/0022-3514.86.1.162

Schmidt, F. L., & Hunter, J. E. (1998). The Validity and Utility of Selection Methods in Personnel Psychology: Practical and Theoretical Implications of 85 Years of Research Findings. Psychological Bulletin, 124(2), 262–274. https://doi.org/10.1037/0033-2909.124.2.262

Schmitt, J. E., Raznahan, A., Clasen, L. S., Wallace, G. L., Pritikin, J. N., Lee, N. R., Giedd, J. N., & Neale, M. C. (2019). The Dynamic Associations between Cortical Thickness and General Intelligence are Genetically Mediated. Cerebral Cortex, 29(11), 4743–4752. https://doi.org/10.1093/cercor/bhz007

Schmitt, N. (2014). Personality and Cognitive Ability as Predictors of Effective Performance at Work. Http://Dx.Doi.Org/10.1146/Annurev-Orgpsych-031413-091255, 1, 45–65. https://doi.org/10.1146/ANNUREV-ORGPSYCH-031413-091255

Schnack, H. G., van Haren, N. E. M., Brouwer, R. M., Evans, A., Durston, S., Boomsma, D. I., Kahn, R. S., & Hulshoff Pol, H. E. (2015). Changes in Thickness and Surface Area of the Human Cortex and Their Relationship with Intelligence. Cerebral Cortex, 25(6), 1608–1617. https://doi.org/10.1093/CERCOR/BHT357

Ségonne, F., Dale, A. M., Busa, E., Glessner, M., Salat, D., Hahn, H. K., & Fischl, B. (2004). A hybrid approach to the skull stripping problem in MRI. NeuroImage, 22(3), 1060–1075. https://doi.org/10.1016/j.neuroimage.2004.03.032

Shaw, P., Greenstein, D., Lerch, J., Clasen, L., Lenroot, R., Gogtay, N., Evans, A., Rapoport, J., & Giedd, J. (2006). Intellectual ability and cortical development in children and adolescents. Nature, 440, 676–679. http://dx.doi.org/10.1038/nature04513

Silver, E., Pulli, E. P., Kataja, E.-L., Kumpulainen, V., Copeland, A., Saukko, E., Saunavaara, J., Merisaari, · Harri Tuire Lähdesmäki, ·, Parkkola, R., Karlsson, L., Karlsson, H., & Tuulari, J. J. (2022). Prenatal and early-life environmental factors, family demographics and cortical brain anatomy in 5-year-olds: an MRI study from FinnBrain Birth Cohort. Brain Imaging and Behavior 2022, 1, 1–13. https://doi.org/10.1007/S11682-022-00679-W

Sled, J. G., Zijdenbos, A. P., & Evans, A. C. (1998). A nonparametric method for automatic correction of intensity nonuniformity in mri data. IEEE Transactions on Medical Imaging, 17(1), 87–97. https://doi.org/10.1109/42.668698

Sølsnes, A. E., Grunewaldt, K. H., Bjuland, K. J., Stavnes, E. M., Bastholm, I. A., Aanes, S., Østgård, H. F., Håberg, A., Løhaugen, G. C. C., Skranes, J., & Rimol, L. M. (2015). Cortical morphometry and IQ in VLBW children without cerebral palsy born in 2003–2007. NeuroImage: Clinical, 8, 193–201. https://doi.org/10.1016/J.NICL.2015.04.004

Sowell, E. R., Thompson, P. M., Leonard, C. M., Welcome, S. E., Kan, E., & Toga, A. W. (2004). Longitudinal Mapping of Cortical Thickness and Brain Growth in Normal Children. Journal of Neuroscience, 24(38), 8223–8231. https://doi.org/10.1523/JNEUROSCI.1798-04.2004

Squeglia, L. M., Jacobus, J., Sorg, S. F., Jernigan, T. L., & Tapert, S. F. (2013). Early Adolescent Cortical Thinning Is Related to Better Neuropsychological Performance. Journal of the International Neuropsychological Society, 19(9), 962–970. https://doi.org/10.1017/S1355617713000878

Strenze, T. (2007). Intelligence and socioeconomic success: A meta-analytic review of longitudinal research. Intelligence, 35(5), 401–426. https://doi.org/10.1016/J.INTELL.2006.09.004

Tadayon, E., Pascual-Leone, A., & Santarnecchi, E. (2020). Differential Contribution of Cortical Thickness, Surface Area, and Gyrification to Fluid and Crystallized Intelligence. Cerebral Cortex, 30(1), 215–225. https://doi.org/10.1093/CERCOR/BHZ082

Thompson, P. M., Cannon, T. D., Narr, K. L., van Erp, T., Poutanen, V.-P., Huttunen, M., Lönnqvist, J., Standertskjöld-Nordenstam, C.-G., Kaprio, J., Khaledy, M., Dail, R., Zoumalan, C. I., & Toga, A. W. (2001). Genetic influences on brain structure. Nature Neuroscience 2001 4:12, 4(12), 1253–1258. https://doi.org/10.1038/NN758

Vargas, T., Damme, K. S. F., & Mittal, V. A. (2020). Neighborhood deprivation, prefrontal morphology and neurocognition in late childhood to early adolescence. NeuroImage, 220, 117086. https://doi.org/10.1016/j.neuroimage.2020.117086

Volkow, N. D., Gordon, J. A., & Freund, M. P. (2021). The Healthy Brain and Child Development Study— Shedding Light on Opioid Exposure, COVID-19, and Health Disparities. JAMA Psychiatry, 78(5), 471–472. https://doi.org/10.1001/JAMAPSYCHIATRY.2020.3803

Volkow, N. D., Koob, G. F., Croyle, R. T., Bianchi, D. W., Gordon, J. A., Koroshetz, W. J., Pérez-Stable, E. J., Riley, W. T., Bloch, M. H., Conway, K., Deeds, B. G., Dowling, G. J., Grant, S., Howlett, K. D., Matochik, J. A., Morgan, G. D., Murray, M. M., Noronha, A., Spong, C. Y., … Weiss, S. R. B. (2018). The conception of the ABCD study: From substance use to a broad NIH collaboration. Developmental Cognitive Neuroscience, 32, 4–7. https://doi.org/10.1016/J.DCN.2017.10.002

Walhovd, K. B., Fjell, A. M., Giedd, J., Dale, A. M., & Brown, T. T. (2016). Through Thick and Thin: a Need to Reconcile Contradictory Results on Trajectories in Human Cortical Development. Cerebral Cortex, 27(2), bhv301. https://doi.org/10.1093/cercor/bhv301

Wallace, G. L., Eric Schmitt, J., Lenroot, R., Viding, E., Ordaz, S., Rosenthal, M. A., Molloy, E. A., Clasen, L. S., Kendler, K. S., Neale, M. C., & Giedd, J. N. (2006). A pediatric twin study of brain morphometry. Journal of Child Psychology and Psychiatry, 47(10), 987–993. https://doi.org/https://doi.org/10.1111/j.1469-7610.2006.01676.x

Wechsler, D. (1967). WPPSI: Wechsler preschool and primary scale of intelligence. New York, N.Y.: Psychological Corporation.

Wechsler, D. (2009). WPPSI-III - Wechsler Preschool And Primary Scale Of Intelligence - Third Edition. Psykologien Kustannus Oy, Helsinki.

Whalley, L. J., & Deary, I. J. (2001). Longitudinal cohort study of childhood IQ and survival up to age 76. British Medical Journal, 322(7290), 819–822. https://doi.org/10.1136/bmj.322.7290.819

Wierenga, L. M., Langen, M., Oranje, B., & Durston, S. (2014). Unique developmental trajectories of cortical thickness and surface area. NeuroImage, 87, 120–126. https://doi.org/10.1016/J.NEUROIMAGE.2013.11.010

Wilke, M., Sohn, J. H., Byars, A. W., & Holland, S. K. (2003). Bright spots: correlations of gray matter volume with IQ in a normal pediatric population. NeuroImage, 20(1), 202–215. https://doi.org/10.1016/S1053-8119(03)00199-X

Winkler, A. M., Kochunov, P., Blangero, J., Almasy, L., Zilles, K., Fox, P. T., Duggirala, R., & Glahn, D. C. (2010). Cortical thickness or grey matter volume? The importance of selecting the phenotype for imaging genetics studies. NeuroImage, 53(3), 1135–1146. https://doi.org/10.1016/J.NEUROIMAGE.2009.12.028

Xuan, D. S., Zhao, X., Liu, Y. C., Xing, Q. N., Shang, H. L., Zhu, P. Y., & Zhang, X. A. (2020). Brain Development in Infants of Mothers With Gestational Diabetes Mellitus: A Diffusion Tensor Imaging Study. Journal of Computer Assisted Tomography, 44(6), 947–952. https://doi.org/10.1097/RCT.0000000000001110

Yokota, S., Takeuchi, H., Hashimoto, T., Hashizume, H., Asano, K., Asano, M., Sassa, Y., Taki, Y., & Kawashima, R. (2015). Individual differences in cognitive performance and brain structure in typically developing children. Developmental Cognitive Neuroscience, 14, 1–7. https://doi.org/10.1016/J.DCN.2015.05.003

