## Supplementary Figure for "Structural brain correlates of non-verbal cognitive ability in 5-year-old children: findings from the FinnBrain Birth Cohort study"

Supplementary Figure 1

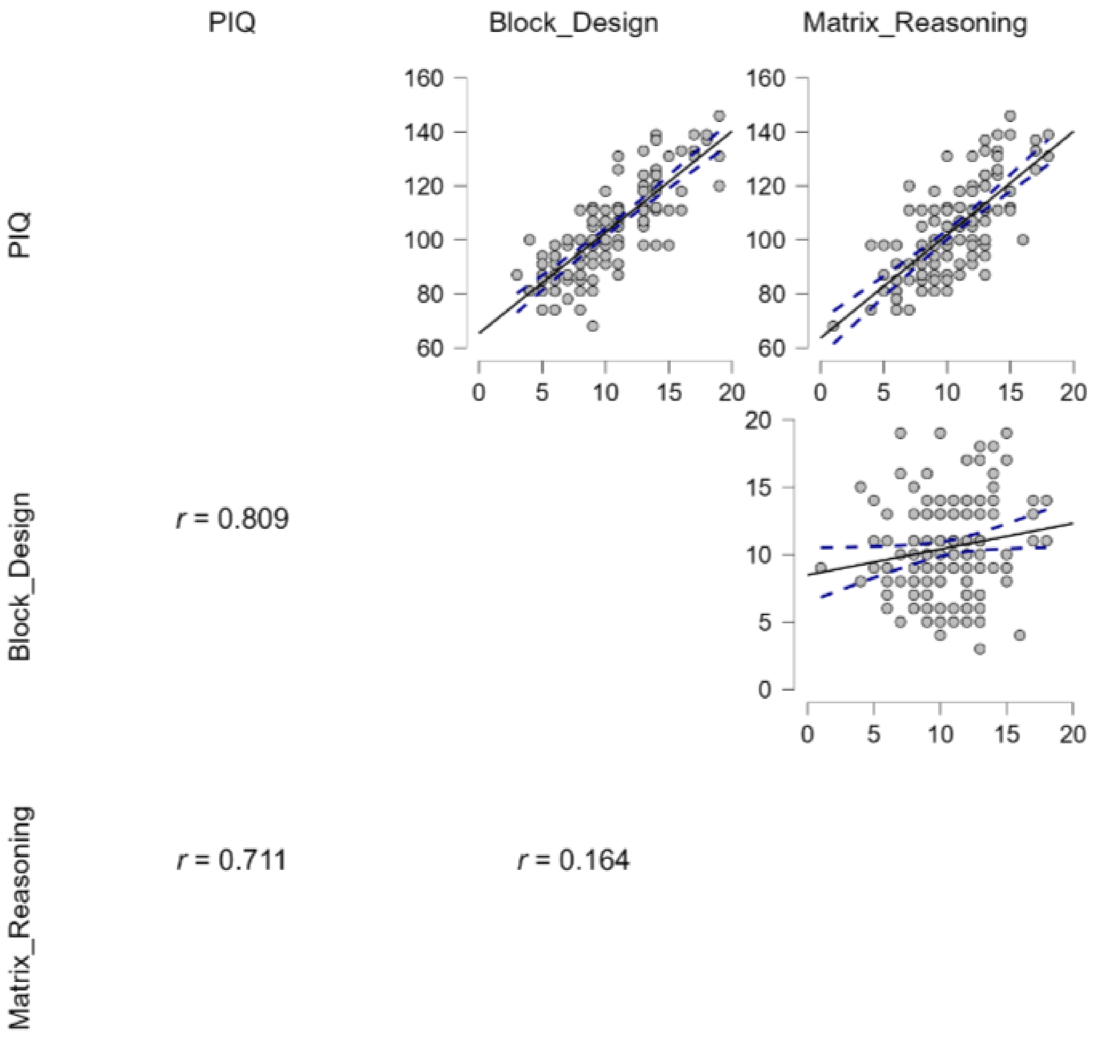


Scatter plots of the Pearson's correlations between performance intelligence quotient (PIQ) and the two subtests it was derived from (Block Design and Matrix Reasoning). All correlations were statistically significant ($p < 0.05$). Dashed line represents the 95% confidence intervals. Normality of distribution was confirmed visually and using the Shapiro-Wilk test (multivariate, $p = 0.143$).

Supplementary Figure 2

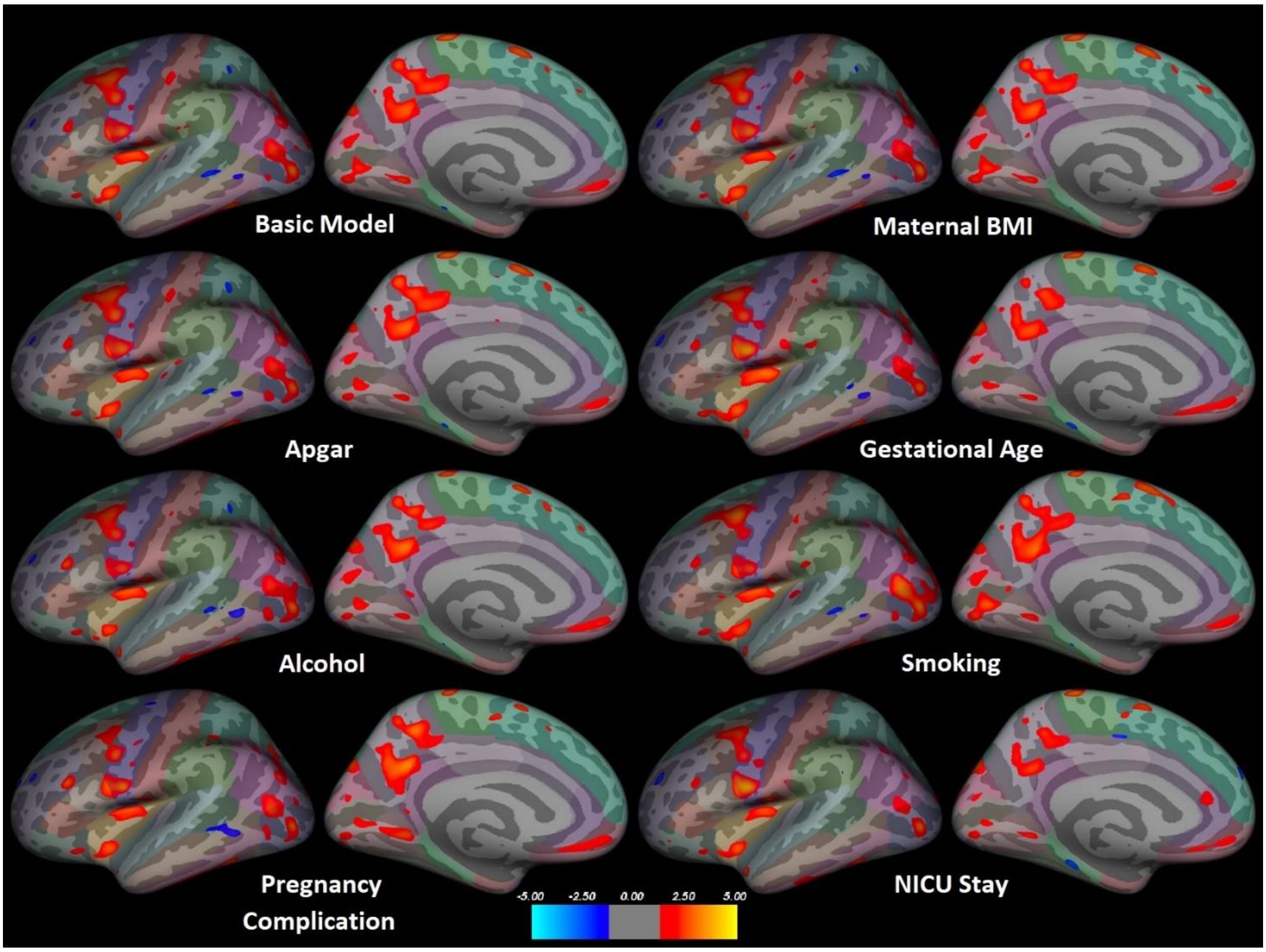


The associations between Performance Intelligence Quotient (PIQ) scores and cortical volumes in the left hemisphere. In top left corner is the model that has been corrected for sex, maternal education level (two classes, university vs. other degree), ponderal index, age at scan (squared), and maternal age at term (the basic model; n = 159). We performed sensitivity analyses, wherein one additional factor was controlled: maternal pre-pregnancy body mass index (BMI; n = 159), or 5 minutes Apgar score (cubed; n = 159). In these two cases, age at scan had to be cubed instead of squared for the purposes of running Qdec. In the other sensitivity analyses, a part of the sample was excluded: children born before gestational week 37 excluded (n = 151), children with prenatal alcohol exposure excluded (alcohol use continued to some degree after learning about the pregnancy, missing data interpreted as no exposure; n = 144), children with prenatal tobacco smoking exposure excluded (smoking at any point during pregnancy, n = 148), children with pregnancy complications excluded (n = 134), and children with neonatal intensive care unit (NICU) stay excluded (n = 136). Cluster color indicates significance as a z-value. Color coding of regions according to the Desikan–Killiany atlas. No correction for multiple comparisons was made.

Supplementary Figure 3

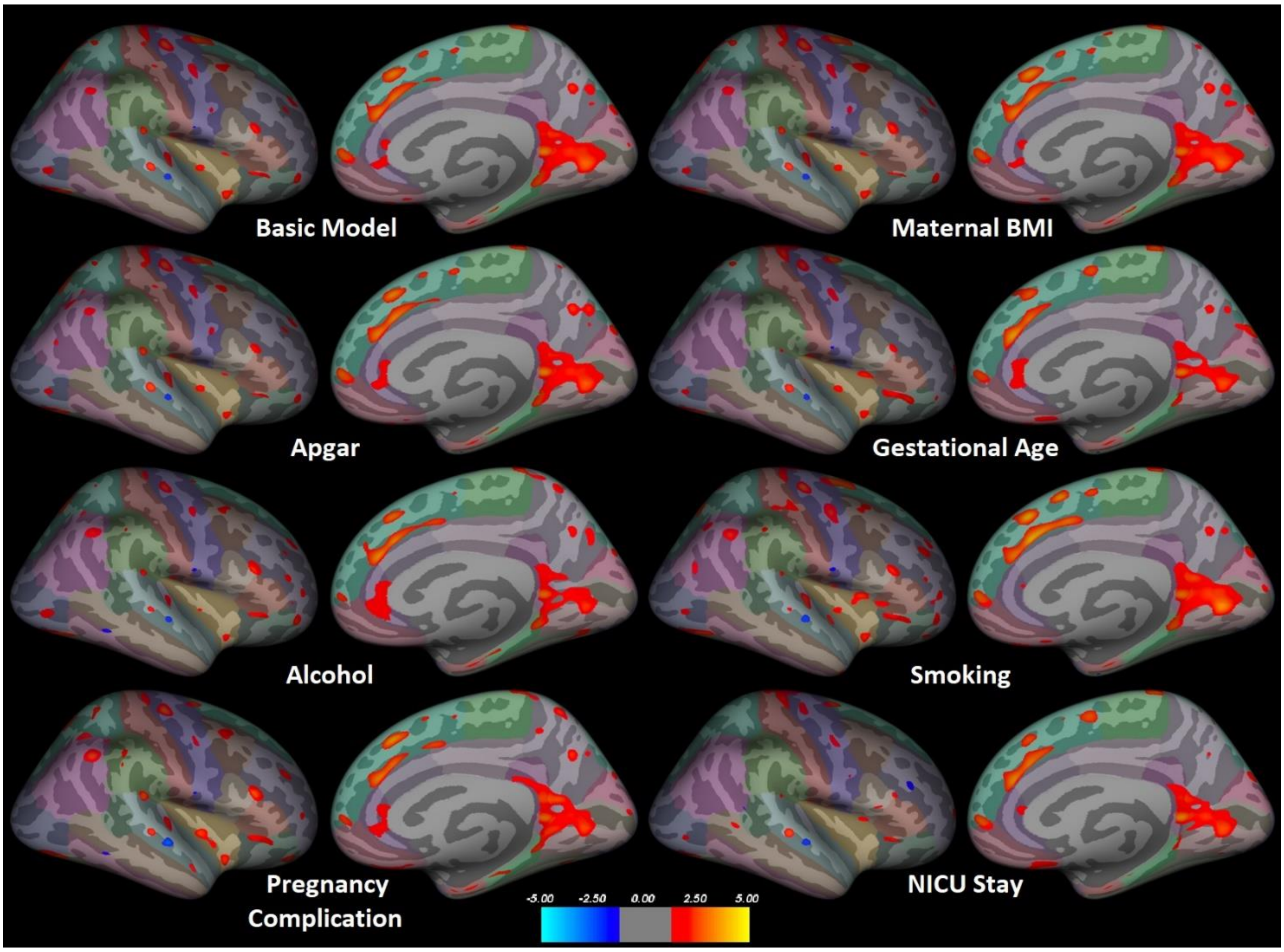


The associations between Performance Intelligence Quotient (PIQ) scores and cortical volumes in the right hemisphere. In top left corner is the model that has been corrected for sex, maternal education level (two classes, university vs. other degree), ponderal index, age at scan (squared), and maternal age at term (the basic model; n = 159). We performed sensitivity analyses, wherein one additional factor was controlled: maternal pre-pregnancy body mass index (BMI; n = 159), or 5 minutes Apgar score (cubed; n = 159). In these two cases, age at scan had to be cubed instead of squared for the purposes of running Qdec. In the other sensitivity analyses, a part of the sample was excluded: children born before gestational week 37 excluded (n = 151), children with prenatal alcohol exposure excluded (alcohol use continued to some degree after learning about the pregnancy, missing data interpreted as no exposure; n = 144), children with prenatal tobacco smoking exposure excluded (smoking at any point during pregnancy, n = 148), children with pregnancy complications excluded (n = 134), and children with neonatal intensive care unit (NICU) stay excluded (n = 136). Cluster color indicates significance as a z-value. Color coding of regions according to the Desikan–Killiany atlas. No correction for multiple comparisons was made.

Supplementary Figure 4

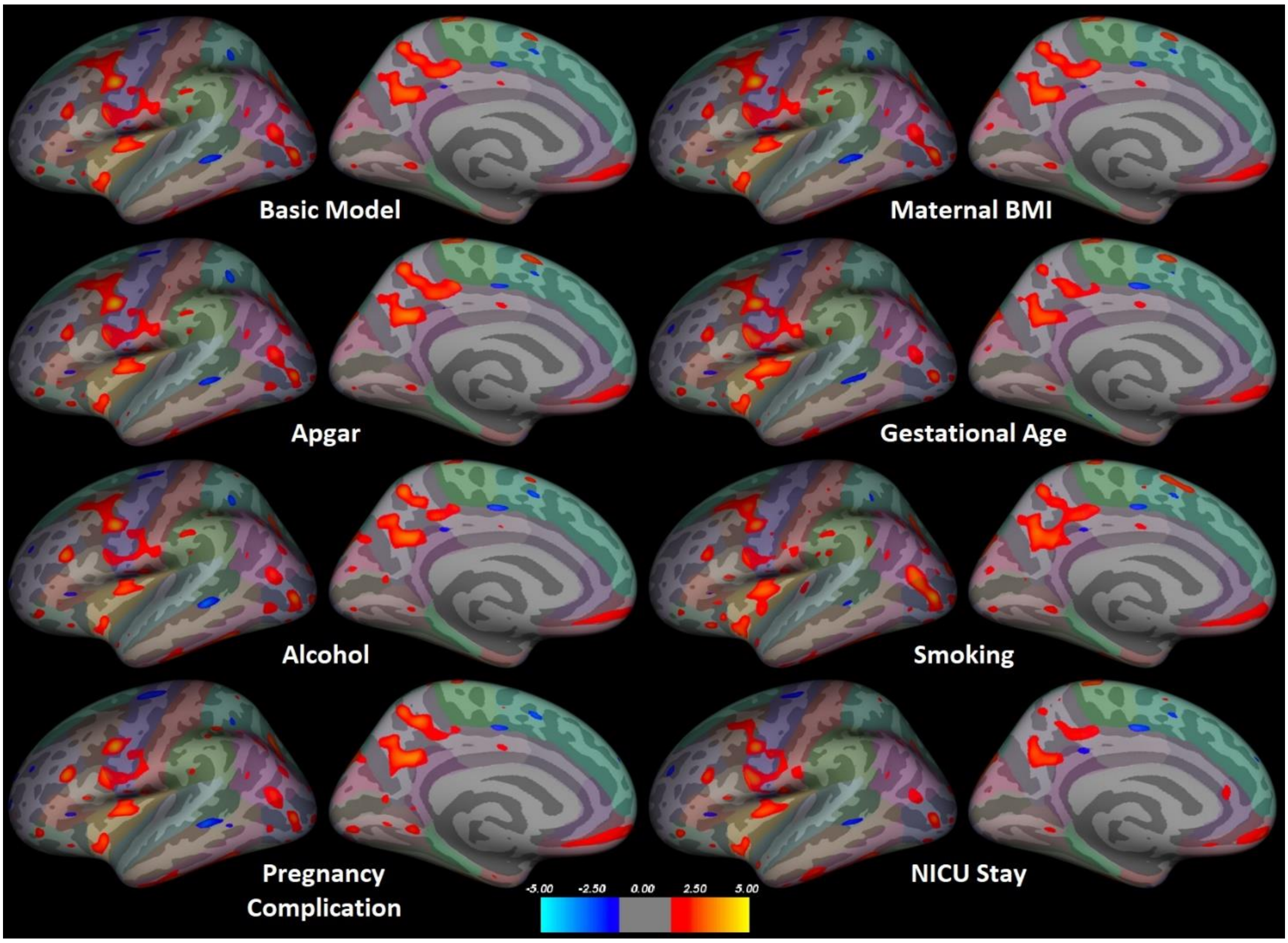


The associations between Performance Intelligence Quotient (PIQ) scores and pial surface area in the left hemisphere. In top left corner is the model that has been corrected for sex, maternal education level (two classes, university vs. other degree), ponderal index, age at scan (squared), and maternal age at term (the basic model; n = 159). We performed sensitivity analyses, wherein one additional factor was controlled: maternal pre-pregnancy body mass index (BMI; n = 159), or 5 minutes Apgar score (cubed; n = 159). In these two cases, age at scan had to be cubed instead of squared for the purposes of running Qdec. In the other sensitivity analyses, a part of the sample was excluded: children born before gestational week 37 excluded (n = 151), children with prenatal alcohol exposure excluded (alcohol use continued to some degree after learning about the pregnancy, missing data interpreted as no exposure; n = 144), children with prenatal tobacco smoking exposure excluded (smoking at any point during pregnancy, n = 148), children with pregnancy complications excluded (n = 134), and children with neonatal intensive care unit (NICU) stay excluded (n = 136). Cluster color indicates significance as a z-value. Color coding of regions according to the Desikan–Killiany atlas. No correction for multiple comparisons was made.

Supplementary Figure 5

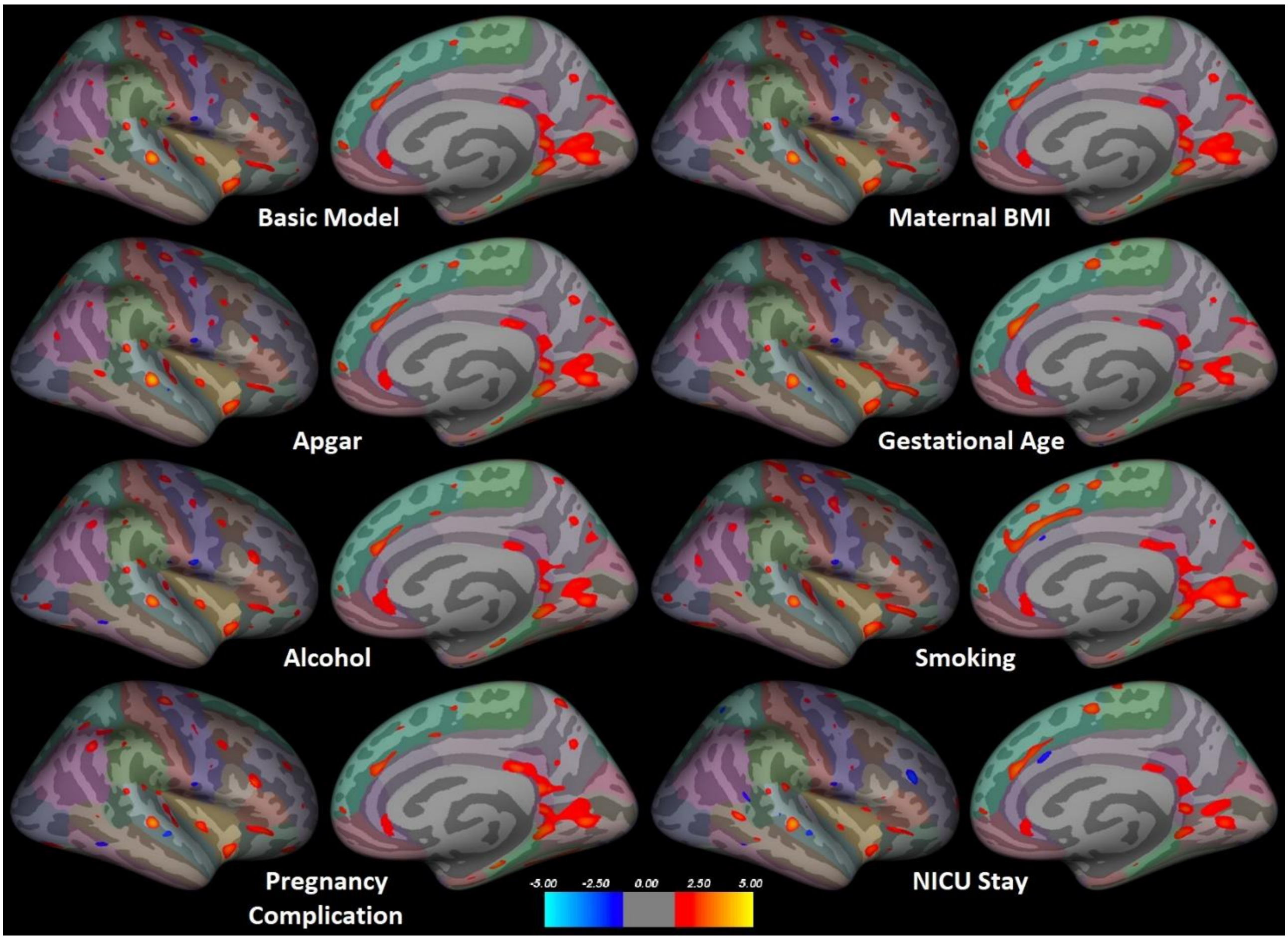


The associations between Performance Intelligence Quotient (PIQ) scores and pial surface area in the right hemisphere. In top left corner is the model that has been corrected for sex, maternal education level (two classes, university vs. other degree), ponderal index, age at scan (squared), and maternal age at term (the basic model; n = 159). We performed sensitivity analyses, wherein one additional factor was controlled: maternal pre-pregnancy body mass index (BMI; n = 159), or 5 minutes Apgar score (cubed; n = 159). In these two cases, age at scan had to be cubed instead of squared for the purposes of running Qdec. In the other sensitivity analyses, a part of the sample was excluded: children born before gestational week 37 excluded (n = 151), children with prenatal alcohol exposure excluded (alcohol use continued to some degree after learning about the pregnancy, missing data interpreted as no exposure; n = 144), children with prenatal tobacco smoking exposure excluded (smoking at any point during pregnancy, n = 148), children with pregnancy complications excluded (n = 134), and children with neonatal intensive care unit (NICU) stay excluded (n = 136). Cluster color indicates significance as a z-value. Color coding of regions according to the Desikan–Killiany atlas. No correction for multiple comparisons was made.

Supplementary Figure 6

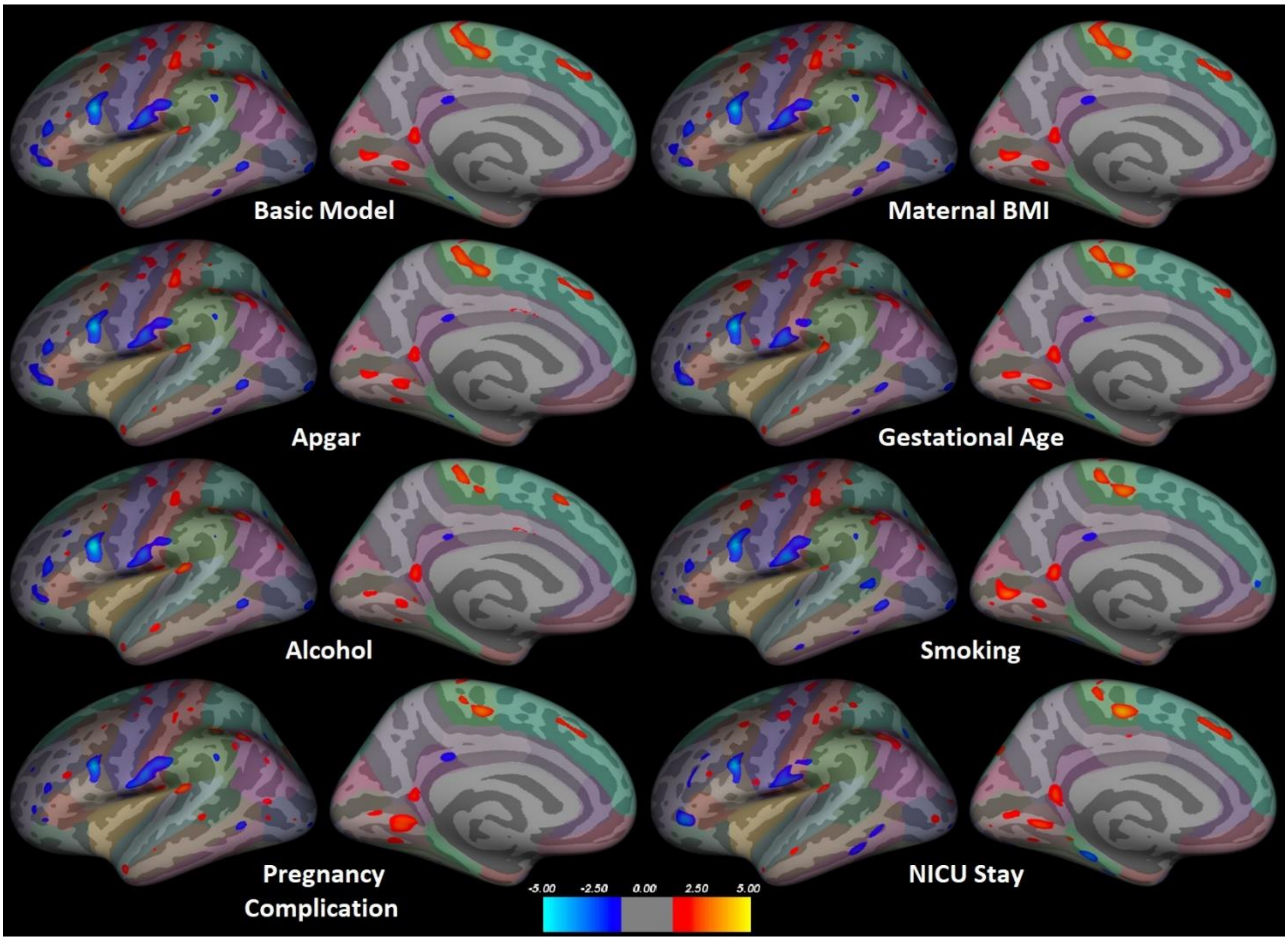


The associations between Performance Intelligence Quotient (PIQ) scores and cortical thickness in the left hemisphere. In top left corner is the model that has been corrected for sex, maternal education level (two classes, university vs. other degree), ponderal index, age at scan (squared), and maternal age at term (the basic model; n = 159). We performed sensitivity analyses, wherein one additional factor was controlled: maternal pre-pregnancy body mass index (BMI; n = 159), or 5 minutes Apgar score (cubed; n = 159). In these two cases, age at scan had to be cubed instead of squared for the purposes of running Qdec. In the other sensitivity analyses, a part of the sample was excluded: children born before gestational week 37 excluded (n = 151), children with prenatal alcohol exposure excluded (alcohol use continued to some degree after learning about the pregnancy, missing data interpreted as no exposure; n = 144), children with prenatal tobacco smoking exposure excluded (smoking at any point during pregnancy, n = 148), children with pregnancy complications excluded (n = 134), and children with neonatal intensive care unit (NICU) stay excluded (n = 136). Cluster color indicates significance as a z-value. Color coding of regions according to the Desikan–Killiany atlas. No correction for multiple comparisons was made.

Supplementary Figure 7

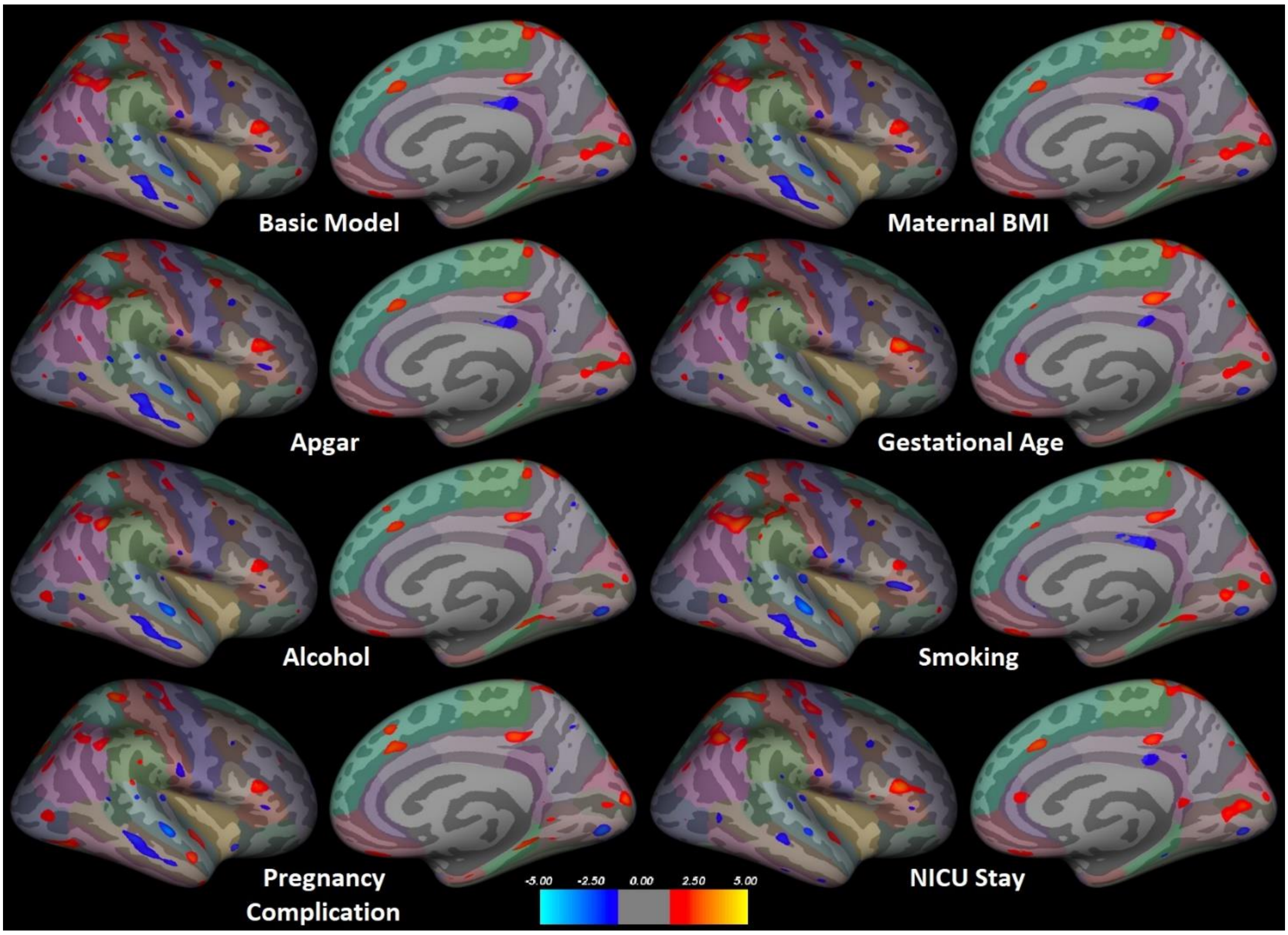


The associations between Performance Intelligence Quotient (PIQ) scores and cortical thickness in the right hemisphere. In top left corner is the model that has been corrected for sex, maternal education level (two classes, university vs. other degree), ponderal index, age at scan (squared), and maternal age at term (the basic model; n = 159). We performed sensitivity analyses, wherein one additional factor was controlled: maternal pre-pregnancy body mass index (BMI; n = 159), or 5 minutes Apgar score (cubed; n = 159). In these two cases, age at scan had to be cubed instead of squared for the purposes of running Qdec. In the other sensitivity analyses, a part of the sample was excluded: children born before gestational week 37 excluded (n = 151), children with prenatal alcohol exposure excluded (alcohol use continued to some degree after learning about the pregnancy, missing data interpreted as no exposure; n = 144), children with prenatal tobacco smoking exposure excluded (smoking at any point during pregnancy, n = 148), children with pregnancy complications excluded (n = 134), and children with neonatal intensive care unit (NICU) stay excluded (n = 136). Cluster color indicates significance as a z-value. Color coding of regions according to the Desikan–Killiany atlas. No correction for multiple comparisons was made.

Supplementary Figure 8

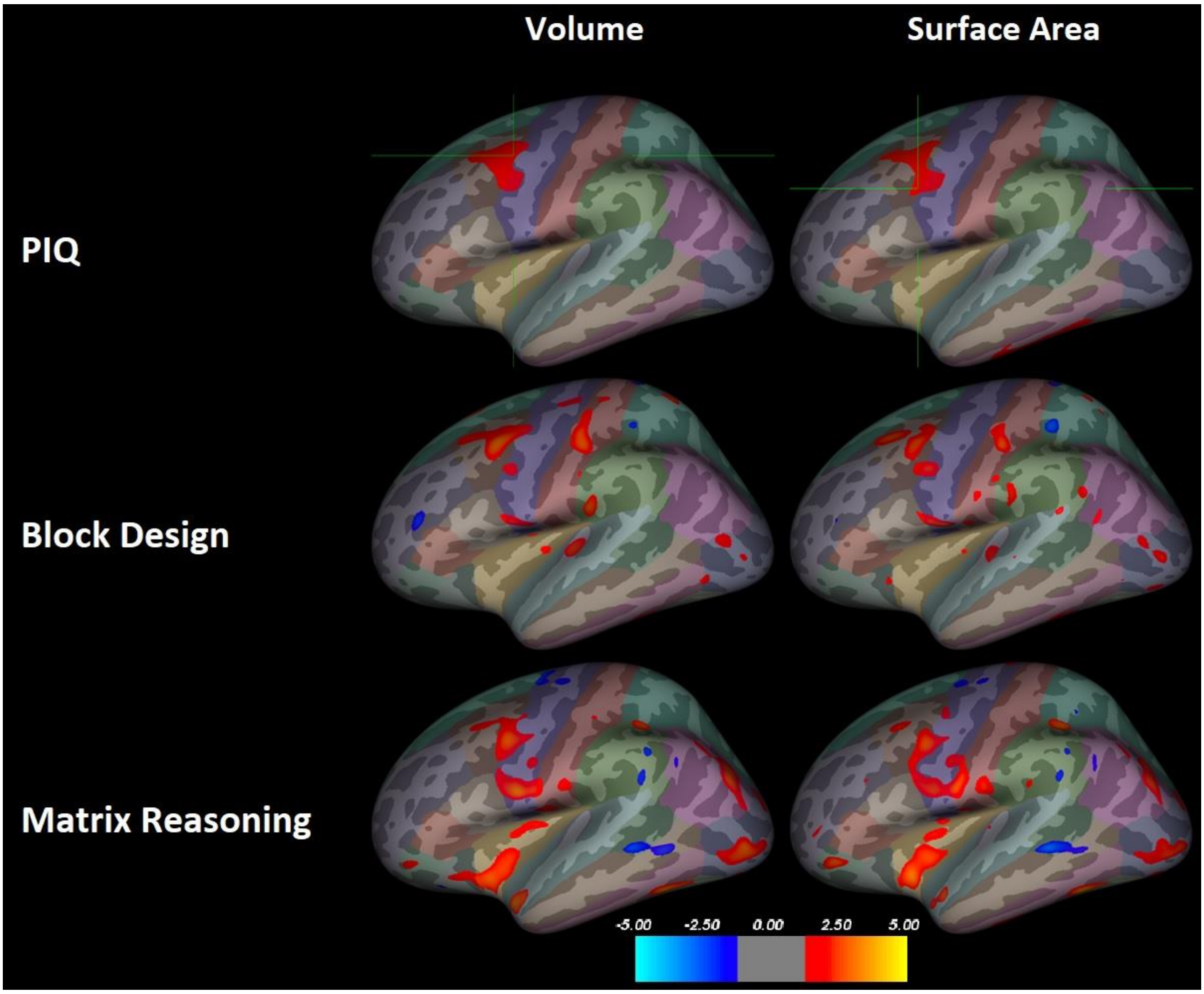


An exploratory analysis showing correlations between performance intelligence quotient (PIQ) and volume in the left caudal middle frontal gyrus, as well as correlations between PIQ and surface area (SA) in the left caudal middle frontal gyrus and the left inferior temporal gyrus (top row). The PIQ results are from Figures 1 and 2 and have been corrected for multiple comparisons using the Monte Carlo simulation. The bottom two rows show the same region for the two subtests without correction for multiple comparisons. In all three clusters, the results don't seem to be strongly driven by either subtest. Cluster color indicates significance as a z-value. The z-value threshold for all images is 1.3. Color coding of regions according to the Desikan–Killiany atlas.

Supplementary Figure 9

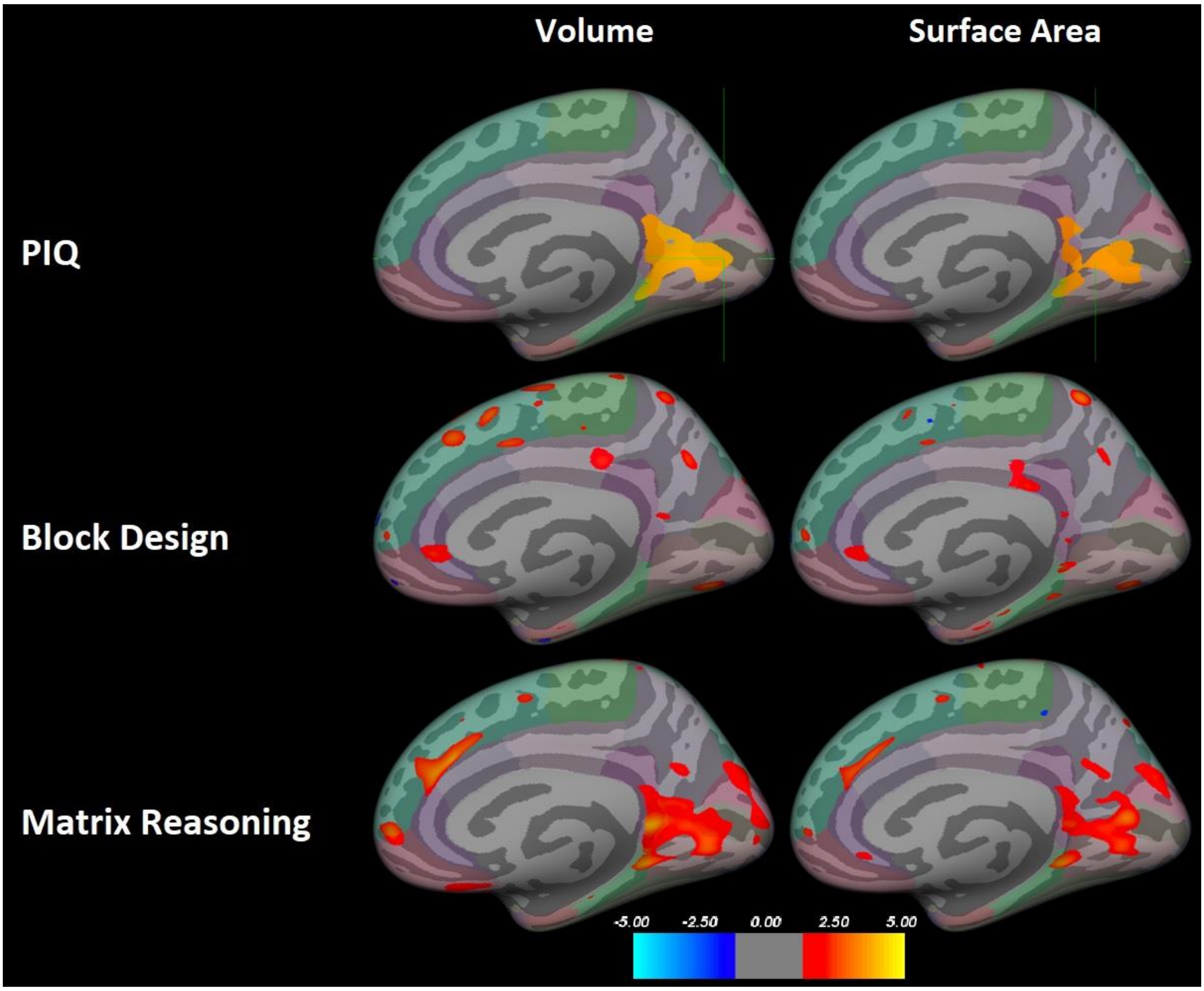


An exploratory analysis showing correlations between performance intelligence quotient (PIQ) and volume/surface area (SA) in the right medial occipital region (top row). The PIQ results are from Figures 1 and 2 and have been corrected for multiple comparisons using the Monte Carlo simulation. The bottom two rows show the same region for the two subtests without correction for multiple comparisons. In both volume and SA, the results seem to be driven by the Matrix Reasoning subtest. Cluster color indicates significance as a z-value. The z-value threshold for all images is 1.3. Color coding of regions according to the Desikan–Killiany atlas.
