## Supplementary Table for "Structural brain correlates of non-verbal cognitive ability in 5-year-old children: findings from the FinnBrain Birth Cohort study"

Partial correlations, controlled for sex, age at scan (in days), ponderal index at scan, maternal education level (university degree vs. any other degree), and maternal age at term
Bonferroni correction for 630 comparisons; target p-value = 0.05/630 = 0.000079.
p < 0.000079
p < 0.05
Abbreviations: ROI = region of interest, PIQ = performance intelligence quotient, df = degree of freedom, lh = left hemisphere, rh = right hemisphere
ROI labels are according to the Desikan-Killiany atlas (https://surfer.nmr.mgh.harvard.edu/fswiki/CorticalParcellation)
ROI exclusions have been made according to the FinnBrain quality control protocol (https://doi.org/10.3389/fnins.2022.874062)

| | | PIQ | Block Design | Matrix Reasoning |
|---|---|---|---|---|
| PIQ | Correlation | 1 | 0.811 | 0.708 |
| | Significance (2-tailed) | . | 2.95E-37 | 1.08E-24 |
| | df | 0 | 152 | 152 |
| Block Design | Correlation | 0.811 | 1 | 0.163 |
| | Significance (2-tailed) | 2.95E-37 | . | 0.043 |
| | df | 152 | 0 | 152 |
| Matrix Reasoning | Correlation | 0.708 | 0.163 | 1 |
| | Significance (2-tailed) | 1.08E-24 | 0.043 | . |
| | df | 152 | 152 | 0 |
| lh_bankssts_volume | Correlation | 0.048 | 0.084 | -0.033 |
| | Significance (2-tailed) | 0.601 | 0.366 | 0.715 |
| | df | 117 | 117 | 122 |
| lh_caudalanteriorcingulate_volume | Correlation | 0.017 | 0.036 | -0.003 |
| | Significance (2-tailed) | 0.847 | 0.687 | 0.971 |
| | df | 125 | 126 | 129 |
| lh_caudalmiddlefrontal_volume | Correlation | 0.138 | 0.122 | 0.09 |
| | Significance (2-tailed) | 0.096 | 0.14 | 0.27 |
| | df | 145 | 146 | 150 |
| lh_cuneus_volume | Correlation | 0.065 | 0.093 | -0.003 |
| | Significance (2-tailed) | 0.495 | 0.33 | 0.971 |
| | df | 109 | 110 | 114 |
| lh_entorhinal_volume | Correlation | 0.063 | 0.08 | 0.031 |
| | Significance (2-tailed) | 0.441 | 0.321 | 0.697 |

| | | | | |
|---|---|---|---|---|
| | df | 152 | 152 | 157 |
| lh_fusiform_volume | Correlation | 0.017 | -0.047 | 0.085 |
| | Significance (2-tailed) | 0.833 | 0.568 | 0.289 |
| | df | 150 | 150 | 155 |
| lh_inferiorparietal_volume | Correlation | 0.105 | 0.058 | 0.118 |
| | Significance (2-tailed) | 0.246 | 0.524 | 0.188 |
| | df | 121 | 121 | 124 |
| lh_inferiortemporal_volume | Correlation | 0.127 | 0.056 | 0.157 |
| | Significance (2-tailed) | 0.151 | 0.531 | 0.071 |
| | df | 127 | 127 | 131 |
| lh_isthmuscingulate_volume | Correlation | 0.099 | 0.074 | 0.102 |
| | Significance (2-tailed) | 0.222 | 0.365 | 0.204 |
| | df | 151 | 152 | 156 |
| lh_lateraloccipital_volume | Correlation | 0.212 | 0.156 | 0.193 |
| | Significance (2-tailed) | 0.016 | 0.078 | 0.027 |
| | df | 126 | 126 | 130 |
| lh_lateralorbitofrontal_volume | Correlation | 0.016 | 0.013 | 0.001 |
| | Significance (2-tailed) | 0.858 | 0.879 | 0.993 |
| | df | 128 | 128 | 133 |
| lh_lingual_volume | Correlation | 0.157 | 0.134 | 0.09 |
| | Significance (2-tailed) | 0.08 | 0.133 | 0.309 |
| | df | 124 | 124 | 128 |
| lh_medialorbitofrontal_volume | Correlation | 0.208 | 0.154 | 0.161 |
| | Significance (2-tailed) | 0.025 | 0.097 | 0.078 |
| | df | 114 | 115 | 119 |
| lh_middletemporal_volume | Correlation | 0.085 | 0.16 | -0.043 |
| | Significance (2-tailed) | 0.408 | 0.119 | 0.671 |
| | df | 94 | 94 | 97 |
| lh_parahippocampal_volume | Correlation | 0.031 | 0.015 | 0.059 |
| | Significance (2-tailed) | 0.7 | 0.853 | 0.462 |
| | df | 151 | 152 | 156 |
| lh_paracentral_volume | Correlation | 0.098 | 0.087 | 0.068 |
| | Significance (2-tailed) | 0.236 | 0.292 | 0.404 |
| | df | 147 | 148 | 152 |
| lh_parsopercularis_volume | Correlation | 0.014 | 0.004 | 0.004 |
| | Significance (2-tailed) | 0.86 | 0.964 | 0.955 |
| | df | 150 | 151 | 155 |
| lh_parsorbitalis_volume | Correlation | 0.063 | 0.039 | 0.054 |
| | Significance (2-tailed) | 0.439 | 0.635 | 0.497 |
| | df | 151 | 152 | 156 |
| lh_parstriangularis_volume | Correlation | 0.004 | 0.004 | -0.005 |
| | Significance (2-tailed) | 0.962 | 0.96 | 0.948 |
| | df | 150 | 151 | 155 |
| lh_pericalcarine_volume | Correlation | 0.172 | 0.187 | 0.063 |
| | Significance (2-tailed) | 0.076 | 0.053 | 0.508 |
| | df | 105 | 105 | 110 |
| lh_postcentral_volu | Correlation | 0.132 | 0.073 | 0.141 |

| | | | | |
|---|---|---|---|---|
| me | Significance (2-tailed) | 0.148 | 0.425 | 0.115 |
| | df | 119 | 120 | 124 |
| lh_posteriorcingulate_volume | Correlation | 0.082 | 0.053 | 0.096 |
| | Significance (2-tailed) | 0.32 | 0.521 | 0.234 |
| | df | 148 | 149 | 153 |
| lh_precentral_volume | Correlation | 0.154 | 0.096 | 0.147 |
| | Significance (2-tailed) | 0.078 | 0.274 | 0.086 |
| | df | 130 | 130 | 135 |
| lh_precuneus_volume | Correlation | 0.26 | 0.218 | 0.168 |
| | Significance (2-tailed) | 0.002 | 0.009 | 0.043 |
| | df | 140 | 141 | 144 |
| lh_rostralanteriorcingulate_volume | Correlation | 0.052 | 0.057 | 0.015 |
| | Significance (2-tailed) | 0.549 | 0.509 | 0.858 |
| | df | 133 | 134 | 138 |
| lh_rostralmiddlefrontal_volume | Correlation | 0.076 | 0.072 | 0.025 |
| | Significance (2-tailed) | 0.355 | 0.379 | 0.756 |
| | df | 148 | 149 | 153 |
| lh_superiorfrontal_volume | Correlation | 0.113 | 0.198 | -0.047 |
| | Significance (2-tailed) | 0.236 | 0.035 | 0.614 |
| | df | 110 | 111 | 114 |
| lh_superiorparietal_volume | Correlation | 0.03 | 0.053 | 0.001 |
| | Significance (2-tailed) | 0.744 | 0.565 | 0.994 |
| | df | 118 | 119 | 122 |
| lh_superiortemporal_volume | Correlation | 0.055 | 0.109 | -0.035 |
| | Significance (2-tailed) | 0.589 | 0.289 | 0.729 |
| | df | 95 | 95 | 98 |
| lh_supramarginal_volume | Correlation | 0.124 | 0.147 | 0.037 |
| | Significance (2-tailed) | 0.214 | 0.139 | 0.71 |
| | df | 101 | 101 | 103 |
| lh_frontalpole_volume | Correlation | 0.051 | 0.036 | 0.037 |
| | Significance (2-tailed) | 0.531 | 0.66 | 0.647 |
| | df | 151 | 152 | 156 |
| lh_temporalpole_volume | Correlation | 0.029 | -0.115 | 0.188 |
| | Significance (2-tailed) | 0.726 | 0.16 | 0.019 |
| | df | 147 | 148 | 152 |
| lh_transversetemporal_volume | Correlation | 0.134 | 0.103 | 0.1 |
| | Significance (2-tailed) | 0.097 | 0.203 | 0.208 |
| | df | 152 | 153 | 157 |
| lh_insula_volume | Correlation | 0.201 | 0.167 | 0.142 |
| | Significance (2-tailed) | 0.025 | 0.063 | 0.107 |
| | df | 122 | 123 | 127 |
| rh_bankssts_volume | Correlation | 0.023 | 0.038 | 0.015 |
| | Significance (2-tailed) | 0.784 | 0.659 | 0.862 |
| | df | 137 | 138 | 142 |
| rh_caudalanteriorcingulate_volume | Correlation | -0.035 | -0.029 | -0.031 |
| | Significance (2-tailed) | 0.686 | 0.742 | 0.715 |
| | df | 132 | 133 | 137 |

| | | | | |
|---|---|---|---|---|
| rh_caudalmiddlefrontal_volume | Correlation | 0.065 | 0.072 | 0.024 |
| | Significance (2-tailed) | 0.435 | 0.388 | 0.767 |
| | df | 144 | 145 | 149 |
| rh_cuneus_volume | Correlation | 0.219 | 0.181 | 0.161 |
| | Significance (2-tailed) | 0.017 | 0.048 | 0.076 |
| | df | 116 | 117 | 121 |
| rh_entorhinal_volume | Correlation | 0.121 | 0.148 | 0.032 |
| | Significance (2-tailed) | 0.136 | 0.067 | 0.688 |
| | df | 151 | 152 | 156 |
| rh_fusiform_volume | Correlation | 0.03 | 0.001 | 0.041 |
| | Significance (2-tailed) | 0.712 | 0.987 | 0.605 |
| | df | 151 | 152 | 156 |
| rh_inferiorparietal_volume | Correlation | 0.016 | 0.062 | -0.054 |
| | Significance (2-tailed) | 0.862 | 0.497 | 0.542 |
| | df | 121 | 121 | 126 |
| rh_inferiortemporal_volume | Correlation | -0.018 | 0.028 | -0.058 |
| | Significance (2-tailed) | 0.849 | 0.769 | 0.534 |
| | df | 113 | 113 | 117 |
| rh_isthmuscingulate_volume | Correlation | 0.123 | 0.047 | 0.158 |
| | Significance (2-tailed) | 0.128 | 0.562 | 0.046 |
| | df | 152 | 153 | 157 |
| rh_lateraloccipital_volume | Correlation | 0.149 | 0.087 | 0.142 |
| | Significance (2-tailed) | 0.078 | 0.308 | 0.09 |
| | df | 138 | 138 | 142 |
| rh_lateralorbitofrontal_volume | Correlation | 0.131 | 0.101 | 0.095 |
| | Significance (2-tailed) | 0.116 | 0.227 | 0.25 |
| | df | 142 | 142 | 146 |
| rh_lingual_volume | Correlation | 0.234 | 0.214 | 0.139 |
| | Significance (2-tailed) | 0.01 | 0.019 | 0.123 |
| | df | 118 | 119 | 123 |
| rh_medialorbitofrontal_volume | Correlation | 0.19 | 0.123 | 0.177 |
| | Significance (2-tailed) | 0.046 | 0.202 | 0.061 |
| | df | 108 | 108 | 111 |
| rh_middletemporal_volume | Correlation | 0.096 | 0.115 | 0.046 |
| | Significance (2-tailed) | 0.37 | 0.278 | 0.66 |
| | df | 88 | 89 | 92 |
| rh_parahippocampal_volume | Correlation | 0.128 | 0.118 | 0.065 |
| | Significance (2-tailed) | 0.113 | 0.144 | 0.415 |
| | df | 152 | 153 | 157 |
| rh_paracentral_volume | Correlation | 0.162 | 0.197 | 0.043 |
| | Significance (2-tailed) | 0.049 | 0.016 | 0.596 |
| | df | 145 | 146 | 149 |
| rh_parsopercularis_volume | Correlation | -0.053 | -0.006 | -0.077 |
| | Significance (2-tailed) | 0.519 | 0.946 | 0.344 |
| | df | 147 | 148 | 151 |
| rh_parsorbitalis_volume | Correlation | 0.179 | 0.146 | 0.126 |
| | Significance (2-tailed) | 0.028 | 0.072 | 0.115 |

| | | | | |
|---|---|---|---|---|
| | df | 150 | 151 | 155 |
| rh_parstriangularis_volume | Correlation | 0.075 | 0.054 | 0.064 |
| | Significance (2-tailed) | 0.362 | 0.511 | 0.426 |
| | df | 149 | 150 | 154 |
| rh_pericalcarine_volume | Correlation | 0.295 | 0.248 | 0.17 |
| | Significance (2-tailed) | 0.002 | 0.008 | 0.067 |
| | df | 110 | 111 | 115 |
| rh_postcentral_volume | Correlation | 0.064 | 0.092 | -0.018 |
| | Significance (2-tailed) | 0.499 | 0.328 | 0.849 |
| | df | 112 | 113 | 115 |
| rh_posteriorcingulate_volume | Correlation | 0.096 | 0.088 | 0.077 |
| | Significance (2-tailed) | 0.241 | 0.28 | 0.337 |
| | df | 149 | 150 | 154 |
| rh_precentral_volume | Correlation | 0.184 | 0.229 | 0.043 |
| | Significance (2-tailed) | 0.042 | 0.011 | 0.633 |
| | df | 121 | 122 | 123 |
| rh_precuneus_volume | Correlation | 0.161 | 0.127 | 0.124 |
| | Significance (2-tailed) | 0.05 | 0.122 | 0.124 |
| | df | 147 | 148 | 152 |
| rh_rostralanteriorcingulate_volume | Correlation | 0.157 | 0.111 | 0.126 |
| | Significance (2-tailed) | 0.054 | 0.174 | 0.116 |
| | df | 149 | 150 | 154 |
| rh_rostralmiddlefrontal_volume | Correlation | 0.139 | 0.128 | 0.072 |
| | Significance (2-tailed) | 0.092 | 0.119 | 0.378 |
| | df | 147 | 148 | 152 |
| rh_superiorfrontal_volume | Correlation | 0.178 | 0.183 | 0.096 |
| | Significance (2-tailed) | 0.046 | 0.04 | 0.276 |
| | df | 124 | 125 | 129 |
| rh_superiorparietal_volume | Correlation | 0.054 | 0.064 | 0.012 |
| | Significance (2-tailed) | 0.569 | 0.5 | 0.901 |
| | df | 112 | 113 | 116 |
| rh_superiortemporal_volume | Correlation | 0.137 | 0.119 | 0.105 |
| | Significance (2-tailed) | 0.157 | 0.219 | 0.269 |
| | df | 106 | 106 | 111 |
| rh_supramarginal_volume | Correlation | 0.175 | 0.145 | 0.122 |
| | Significance (2-tailed) | 0.067 | 0.13 | 0.194 |
| | df | 108 | 108 | 113 |
| rh_frontalpole_volume | Correlation | 0.025 | -0.016 | 0.058 |
| | Significance (2-tailed) | 0.761 | 0.841 | 0.466 |
| | df | 151 | 152 | 156 |
| rh_temporalpole_volume | Correlation | 0.04 | 0.056 | 0.009 |
| | Significance (2-tailed) | 0.631 | 0.496 | 0.908 |
| | df | 148 | 149 | 153 |
| rh_transversetemporal_volume | Correlation | 0.153 | 0.178 | 0.036 |
| | Significance (2-tailed) | 0.059 | 0.027 | 0.648 |
| | df | 152 | 153 | 157 |
| rh_insula_volume | Correlation | 0.237 | 0.18 | 0.188 |

| | | | | |
|---|---|---|---|---|
| | Significance (2-tailed) | 0.009 | 0.049 | 0.038 |
| | df | 117 | 118 | 121 |
| BrainSegVolNotVent | Correlation | 0.188 | 0.181 | 0.099 |
| | Significance (2-tailed) | 0.019 | 0.024 | 0.215 |
| | df | 152 | 153 | 157 |
| eTIV | Correlation | 0.144 | 0.16 | 0.06 |
| | Significance (2-tailed) | 0.074 | 0.047 | 0.453 |
| | df | 152 | 153 | 157 |
| lh_bankssts_area | Correlation | 0.054 | 0.081 | -0.022 |
| | Significance (2-tailed) | 0.56 | 0.382 | 0.807 |
| | df | 117 | 117 | 122 |
| lh_caudalanteriorcingulate_area | Correlation | 0.029 | 0.041 | 0.016 |
| | Significance (2-tailed) | 0.748 | 0.642 | 0.86 |
| | df | 125 | 126 | 129 |
| lh_caudalmiddlefrontal_area | Correlation | 0.094 | 0.084 | 0.069 |
| | Significance (2-tailed) | 0.255 | 0.307 | 0.401 |
| | df | 145 | 146 | 150 |
| lh_cuneus_area | Correlation | 0.098 | 0.094 | 0.052 |
| | Significance (2-tailed) | 0.308 | 0.324 | 0.578 |
| | df | 109 | 110 | 114 |
| lh_entorhinal_area | Correlation | 0.027 | 0.032 | 0.021 |
| | Significance (2-tailed) | 0.743 | 0.691 | 0.796 |
| | df | 152 | 152 | 157 |
| lh_fusiform_area | Correlation | 0.066 | 0.018 | 0.084 |
| | Significance (2-tailed) | 0.418 | 0.822 | 0.297 |
| | df | 150 | 150 | 155 |
| lh_inferiorparietal_area | Correlation | 0.067 | 0.027 | 0.079 |
| | Significance (2-tailed) | 0.464 | 0.768 | 0.381 |
| | df | 121 | 121 | 124 |
| lh_inferiortemporal_area | Correlation | 0.191 | 0.104 | 0.2 |
| | Significance (2-tailed) | 0.03 | 0.239 | 0.021 |
| | df | 127 | 127 | 131 |
| lh_isthmuscingulate_area | Correlation | 0.099 | 0.076 | 0.086 |
| | Significance (2-tailed) | 0.222 | 0.351 | 0.283 |
| | df | 151 | 152 | 156 |
| lh_lateraloccipital_area | Correlation | 0.161 | 0.097 | 0.164 |
| | Significance (2-tailed) | 0.069 | 0.275 | 0.06 |
| | df | 126 | 126 | 130 |
| lh_lateralorbitofrontal_area | Correlation | 0.051 | 0.037 | 0.023 |
| | Significance (2-tailed) | 0.562 | 0.677 | 0.794 |
| | df | 128 | 128 | 133 |
| lh_lingual_area | Correlation | 0.123 | 0.094 | 0.085 |
| | Significance (2-tailed) | 0.171 | 0.294 | 0.335 |
| | df | 124 | 124 | 128 |
| lh_medialorbitofrontal_area | Correlation | 0.256 | 0.163 | 0.232 |
| | Significance (2-tailed) | 0.006 | 0.079 | 0.01 |
| | df | 114 | 115 | 119 |

| | | | | |
|---|---|---|---|---|
| lh_middletemporal_area | Correlation | 0.046 | 0.114 | -0.069 |
| | Significance (2-tailed) | 0.66 | 0.27 | 0.497 |
| | df | 94 | 94 | 97 |
| lh_parahippocampal_area | Correlation | 0.002 | 0.077 | -0.098 |
| | Significance (2-tailed) | 0.978 | 0.341 | 0.221 |
| | df | 151 | 152 | 156 |
| lh_paracentral_area | Correlation | 0.014 | 0.011 | 0.021 |
| | Significance (2-tailed) | 0.864 | 0.898 | 0.795 |
| | df | 147 | 148 | 152 |
| lh_parsopercularis_area | Correlation | 0.044 | 0.03 | 0.018 |
| | Significance (2-tailed) | 0.586 | 0.714 | 0.827 |
| | df | 150 | 151 | 155 |
| lh_parsorbitalis_area | Correlation | 0.091 | 0.09 | 0.027 |
| | Significance (2-tailed) | 0.262 | 0.265 | 0.734 |
| | df | 151 | 152 | 156 |
| lh_parstriangularis_area | Correlation | 0.056 | 0.064 | 0 |
| | Significance (2-tailed) | 0.491 | 0.434 | 0.997 |
| | df | 150 | 151 | 155 |
| lh_pericalcarine_area | Correlation | 0.121 | 0.142 | 0.028 |
| | Significance (2-tailed) | 0.213 | 0.144 | 0.768 |
| | df | 105 | 105 | 110 |
| lh_postcentral_area | Correlation | 0.049 | -0.002 | 0.1 |
| | Significance (2-tailed) | 0.594 | 0.98 | 0.264 |
| | df | 119 | 120 | 124 |
| lh_posteriorcingulate_area | Correlation | 0.088 | 0.092 | 0.048 |
| | Significance (2-tailed) | 0.284 | 0.261 | 0.554 |
| | df | 148 | 149 | 153 |
| lh_precentral_area | Correlation | 0.076 | 0.031 | 0.101 |
| | Significance (2-tailed) | 0.385 | 0.722 | 0.242 |
| | df | 130 | 130 | 135 |
| lh_precuneus_area | Correlation | 0.217 | 0.17 | 0.146 |
| | Significance (2-tailed) | 0.009 | 0.043 | 0.078 |
| | df | 140 | 141 | 144 |
| lh_rostralanteriorcingulate_area | Correlation | 0.101 | 0.127 | 0.021 |
| | Significance (2-tailed) | 0.244 | 0.14 | 0.805 |
| | df | 133 | 134 | 138 |
| lh_rostralmiddlefrontal_area | Correlation | 0.068 | 0.069 | 0.018 |
| | Significance (2-tailed) | 0.405 | 0.399 | 0.821 |
| | df | 148 | 149 | 153 |
| lh_superiorfrontal_area | Correlation | 0.007 | 0.092 | -0.089 |
| | Significance (2-tailed) | 0.942 | 0.331 | 0.342 |
| | df | 110 | 111 | 114 |
| lh_superiorparietal_area | Correlation | -0.043 | -0.037 | -0.008 |
| | Significance (2-tailed) | 0.644 | 0.687 | 0.927 |
| | df | 118 | 119 | 122 |
| lh_superiortemporal_area | Correlation | 0.037 | 0.072 | -0.028 |
| | Significance (2-tailed) | 0.716 | 0.481 | 0.781 |

| | | | | |
|---|---|---|---|---|
| | df | 95 | 95 | 98 |
| lh_supramarginal_area | Correlation | 0.083 | 0.097 | 0.023 |
| | Significance (2-tailed) | 0.405 | 0.33 | 0.819 |
| | df | 101 | 101 | 103 |
| lh_frontalpole_area | Correlation | 0.053 | 0.009 | 0.08 |
| | Significance (2-tailed) | 0.516 | 0.916 | 0.32 |
| | df | 151 | 152 | 156 |
| lh_temporalpole_area | Correlation | 0.059 | -0.021 | 0.137 |
| | Significance (2-tailed) | 0.478 | 0.796 | 0.09 |
| | df | 147 | 148 | 152 |
| lh_transversetemporal_area | Correlation | 0.127 | 0.039 | 0.17 |
| | Significance (2-tailed) | 0.117 | 0.63 | 0.032 |
| | df | 152 | 153 | 157 |
| lh_insula_area | Correlation | 0.263 | 0.197 | 0.2 |
| | Significance (2-tailed) | 0.003 | 0.028 | 0.023 |
| | df | 122 | 123 | 127 |
| rh_bankssts_area | Correlation | 0.017 | 0.031 | -0.006 |
| | Significance (2-tailed) | 0.845 | 0.719 | 0.939 |
| | df | 137 | 138 | 142 |
| rh_caudalanteriorcingulate_area | Correlation | 0.035 | 0.038 | 0.008 |
| | Significance (2-tailed) | 0.688 | 0.659 | 0.925 |
| | df | 132 | 133 | 137 |
| rh_caudalmiddlefrontal_area | Correlation | 0.002 | 0.04 | -0.036 |
| | Significance (2-tailed) | 0.984 | 0.633 | 0.658 |
| | df | 144 | 145 | 149 |
| rh_cuneus_area | Correlation | 0.174 | 0.155 | 0.116 |
| | Significance (2-tailed) | 0.059 | 0.092 | 0.2 |
| | df | 116 | 117 | 121 |
| rh_entorhinal_area | Correlation | 0.051 | 0.088 | -0.007 |
| | Significance (2-tailed) | 0.53 | 0.278 | 0.928 |
| | df | 151 | 152 | 156 |
| rh_fusiform_area | Correlation | 0.055 | 0.026 | 0.048 |
| | Significance (2-tailed) | 0.497 | 0.752 | 0.55 |
| | df | 151 | 152 | 156 |
| rh_inferiorparietal_area | Correlation | -0.023 | 0.02 | -0.074 |
| | Significance (2-tailed) | 0.805 | 0.824 | 0.408 |
| | df | 121 | 121 | 126 |
| rh_inferiortemporal_area | Correlation | -0.015 | -0.034 | 0.02 |
| | Significance (2-tailed) | 0.872 | 0.717 | 0.831 |
| | df | 113 | 113 | 117 |
| rh_isthmuscingulate_area | Correlation | 0.146 | 0.084 | 0.152 |
| | Significance (2-tailed) | 0.07 | 0.3 | 0.056 |
| | df | 152 | 153 | 157 |
| rh_lateraloccipital_area | Correlation | 0.094 | 0.019 | 0.134 |
| | Significance (2-tailed) | 0.267 | 0.822 | 0.109 |
| | df | 138 | 138 | 142 |
| rh_lateralorbitofront | Correlation | 0.2 | 0.167 | 0.125 |

| | | | | |
|---|---|---|---|---|
| al_area | Significance (2-tailed) | 0.016 | 0.045 | 0.13 |
| | df | 142 | 142 | 146 |
| rh_lingual_area | Correlation | 0.235 | 0.233 | 0.117 |
| | Significance (2-tailed) | 0.01 | 0.01 | 0.194 |
| | df | 118 | 119 | 123 |
| rh_medialorbitofrontal_area | Correlation | 0.13 | 0.118 | 0.07 |
| | Significance (2-tailed) | 0.176 | 0.219 | 0.464 |
| | df | 108 | 108 | 111 |
| rh_middletemporal_area | Correlation | 0.071 | 0.095 | 0.016 |
| | Significance (2-tailed) | 0.505 | 0.371 | 0.875 |
| | df | 88 | 89 | 92 |
| rh_parahippocampal_area | Correlation | 0.037 | 0.087 | -0.067 |
| | Significance (2-tailed) | 0.649 | 0.281 | 0.405 |
| | df | 152 | 153 | 157 |
| rh_paracentral_area | Correlation | 0.123 | 0.145 | 0.042 |
| | Significance (2-tailed) | 0.137 | 0.08 | 0.611 |
| | df | 145 | 146 | 149 |
| rh_parsopercularis_area | Correlation | -0.049 | -0.025 | -0.052 |
| | Significance (2-tailed) | 0.555 | 0.763 | 0.519 |
| | df | 147 | 148 | 151 |
| rh_parsorbitalis_area | Correlation | 0.174 | 0.153 | 0.086 |
| | Significance (2-tailed) | 0.032 | 0.059 | 0.282 |
| | df | 150 | 151 | 155 |
| rh_parstriangularis_area | Correlation | 0.062 | 0.045 | 0.04 |
| | Significance (2-tailed) | 0.451 | 0.578 | 0.616 |
| | df | 149 | 150 | 154 |
| rh_pericalcarine_area | Correlation | 0.208 | 0.242 | 0.063 |
| | Significance (2-tailed) | 0.028 | 0.01 | 0.501 |
| | df | 110 | 111 | 115 |
| rh_postcentral_area | Correlation | -0.021 | -0.01 | -0.031 |
| | Significance (2-tailed) | 0.828 | 0.917 | 0.742 |
| | df | 112 | 113 | 115 |
| rh_posteriorcingulate_area | Correlation | 0.121 | 0.123 | 0.067 |
| | Significance (2-tailed) | 0.14 | 0.13 | 0.408 |
| | df | 149 | 150 | 154 |
| rh_precentral_area | Correlation | 0.06 | 0.116 | -0.038 |
| | Significance (2-tailed) | 0.513 | 0.2 | 0.671 |
| | df | 121 | 122 | 123 |
| rh_precuneus_area | Correlation | 0.119 | 0.072 | 0.118 |
| | Significance (2-tailed) | 0.149 | 0.384 | 0.146 |
| | df | 147 | 148 | 152 |
| rh_rostralanteriorcingulate_area | Correlation | 0.147 | 0.126 | 0.088 |
| | Significance (2-tailed) | 0.072 | 0.122 | 0.274 |
| | df | 149 | 150 | 154 |
| rh_rostralmiddlefrontal_area | Correlation | 0.117 | 0.122 | 0.041 |
| | Significance (2-tailed) | 0.154 | 0.136 | 0.612 |
| | df | 147 | 148 | 152 |

| | | | | |
|---|---|---|---|---|
| rh_superiorfrontal_area | Correlation | 0.101 | 0.127 | 0.024 |
| | Significance (2-tailed) | 0.261 | 0.156 | 0.782 |
| | df | 124 | 125 | 129 |
| rh_superiorparietal_area | Correlation | -0.054 | -0.034 | -0.047 |
| | Significance (2-tailed) | 0.572 | 0.719 | 0.611 |
| | df | 112 | 113 | 116 |
| rh_superiortemporal_area | Correlation | 0.147 | 0.132 | 0.099 |
| | Significance (2-tailed) | 0.129 | 0.174 | 0.299 |
| | df | 106 | 106 | 111 |
| rh_supramarginal_area | Correlation | 0.147 | 0.112 | 0.105 |
| | Significance (2-tailed) | 0.124 | 0.245 | 0.264 |
| | df | 108 | 108 | 113 |
| rh_frontalpole_area | Correlation | 0.056 | 0.013 | 0.071 |
| | Significance (2-tailed) | 0.493 | 0.874 | 0.377 |
| | df | 151 | 152 | 156 |
| rh_temporalpole_area | Correlation | 0.068 | 0.063 | 0.057 |
| | Significance (2-tailed) | 0.409 | 0.445 | 0.484 |
| | df | 148 | 149 | 153 |
| rh_transversetemporal_area | Correlation | 0.157 | 0.119 | 0.11 |
| | Significance (2-tailed) | 0.051 | 0.139 | 0.167 |
| | df | 152 | 153 | 157 |
| rh_insula_area | Correlation | 0.29 | 0.272 | 0.168 |
| | Significance (2-tailed) | 0.001 | 0.003 | 0.063 |
| | df | 117 | 118 | 121 |
| lh_surface_area | Correlation | 0.151 | 0.12 | 0.106 |
| | Significance (2-tailed) | 0.061 | 0.139 | 0.182 |
| | df | 152 | 153 | 157 |
| rh_surface_area | Correlation | 0.135 | 0.124 | 0.074 |
| | Significance (2-tailed) | 0.096 | 0.124 | 0.356 |
| | df | 152 | 153 | 157 |
| lh_bankssts_thickness | Correlation | -0.007 | 0.02 | -0.033 |
| | Significance (2-tailed) | 0.939 | 0.828 | 0.713 |
| | df | 117 | 117 | 122 |
| lh_caudalanteriorcingulate_thickness | Correlation | 0.038 | 0.03 | 0.01 |
| | Significance (2-tailed) | 0.672 | 0.737 | 0.912 |
| | df | 125 | 126 | 129 |
| lh_caudalmiddlefrontal_thickness | Correlation | 0.066 | 0.049 | 0.028 |
| | Significance (2-tailed) | 0.426 | 0.552 | 0.733 |
| | df | 145 | 146 | 150 |
| lh_cuneus_thickness | Correlation | -0.047 | 0.027 | -0.121 |
| | Significance (2-tailed) | 0.627 | 0.776 | 0.194 |
| | df | 109 | 110 | 114 |
| lh_entorhinal_thickness | Correlation | 0.061 | 0.05 | 0.051 |
| | Significance (2-tailed) | 0.454 | 0.54 | 0.521 |
| | df | 152 | 152 | 157 |
| lh_fusiform_thickness | Correlation | -0.036 | -0.057 | 0.02 |
| | Significance (2-tailed) | 0.66 | 0.487 | 0.806 |

| | | | | |
|---|---|---|---|---|
| | df | 150 | 150 | 155 |
| lh_inferiorparietal_thickness | Correlation | 0.086 | 0.069 | 0.086 |
| | Significance (2-tailed) | 0.344 | 0.446 | 0.338 |
| | df | 121 | 121 | 124 |
| lh_inferiortemporal_thickness | Correlation | -0.097 | -0.1 | -0.025 |
| | Significance (2-tailed) | 0.275 | 0.259 | 0.772 |
| | df | 127 | 127 | 131 |
| lh_isthmuscingulate_thickness | Correlation | 0.076 | 0.062 | 0.072 |
| | Significance (2-tailed) | 0.353 | 0.443 | 0.368 |
| | df | 151 | 152 | 156 |
| lh_lateraloccipital_thickness | Correlation | 0.139 | 0.149 | 0.082 |
| | Significance (2-tailed) | 0.117 | 0.093 | 0.348 |
| | df | 126 | 126 | 130 |
| lh_lateralorbitofrontal_thickness | Correlation | -0.024 | -0.042 | 0.024 |
| | Significance (2-tailed) | 0.784 | 0.637 | 0.78 |
| | df | 128 | 128 | 133 |
| lh_lingual_thickness | Correlation | 0.106 | 0.106 | 0.038 |
| | Significance (2-tailed) | 0.236 | 0.236 | 0.671 |
| | df | 124 | 124 | 128 |
| lh_medialorbitofrontal_thickness | Correlation | -0.108 | -0.044 | -0.131 |
| | Significance (2-tailed) | 0.247 | 0.634 | 0.151 |
| | df | 114 | 115 | 119 |
| lh_middletemporal_thickness | Correlation | 0.107 | 0.108 | 0.092 |
| | Significance (2-tailed) | 0.299 | 0.297 | 0.366 |
| | df | 94 | 94 | 97 |
| lh_parahippocampal_thickness | Correlation | -0.015 | -0.074 | 0.102 |
| | Significance (2-tailed) | 0.85 | 0.362 | 0.201 |
| | df | 151 | 152 | 156 |
| lh_paracentral_thickness | Correlation | 0.21 | 0.223 | 0.069 |
| | Significance (2-tailed) | 0.01 | 0.006 | 0.392 |
| | df | 147 | 148 | 152 |
| lh_parsopercularis_thickness | Correlation | -0.076 | -0.069 | -0.026 |
| | Significance (2-tailed) | 0.353 | 0.399 | 0.748 |
| | df | 150 | 151 | 155 |
| lh_parsorbitalis_thickness | Correlation | -0.039 | -0.017 | -0.031 |
| | Significance (2-tailed) | 0.63 | 0.837 | 0.703 |
| | df | 151 | 152 | 156 |
| lh_parstriangularis_thickness | Correlation | -0.071 | -0.063 | -0.015 |
| | Significance (2-tailed) | 0.385 | 0.437 | 0.851 |
| | df | 150 | 151 | 155 |
| lh_pericalcarine_thickness | Correlation | 0.171 | 0.177 | 0.079 |
| | Significance (2-tailed) | 0.079 | 0.068 | 0.41 |
| | df | 105 | 105 | 110 |
| lh_postcentral_thickness | Correlation | 0.135 | 0.134 | 0.049 |
| | Significance (2-tailed) | 0.14 | 0.141 | 0.582 |
| | df | 119 | 120 | 124 |
| lh_posteriorcingulate | Correlation | 0.037 | -0.026 | 0.111 |

| | | | | |
|---|---|---|---|---|
| _thickness | Significance (2-tailed) | 0.652 | 0.756 | 0.168 |
| | df | 148 | 149 | 153 |
| lh_precentral_thickness | Correlation | 0.156 | 0.159 | 0.062 |
| | Significance (2-tailed) | 0.074 | 0.068 | 0.471 |
| | df | 130 | 130 | 135 |
| lh_precuneus_thickness | Correlation | 0.051 | 0.076 | -0.002 |
| | Significance (2-tailed) | 0.544 | 0.366 | 0.98 |
| | df | 140 | 141 | 144 |
| lh_rostralanteriorcingulate_thickness | Correlation | -0.031 | -0.045 | -0.005 |
| | Significance (2-tailed) | 0.725 | 0.6 | 0.954 |
| | df | 133 | 134 | 138 |
| lh_rostralmiddlefrontal_thickness | Correlation | -0.017 | -0.01 | -0.014 |
| | Significance (2-tailed) | 0.838 | 0.9 | 0.866 |
| | df | 148 | 149 | 153 |
| lh_superiorfrontal_thickness | Correlation | 0.172 | 0.159 | 0.074 |
| | Significance (2-tailed) | 0.07 | 0.093 | 0.43 |
| | df | 110 | 111 | 114 |
| lh_superiorparietal_thickness | Correlation | 0.165 | 0.197 | 0.029 |
| | Significance (2-tailed) | 0.071 | 0.03 | 0.751 |
| | df | 118 | 119 | 122 |
| lh_superiortemporal_thickness | Correlation | -0.006 | 0.026 | -0.032 |
| | Significance (2-tailed) | 0.95 | 0.8 | 0.749 |
| | df | 95 | 95 | 98 |
| lh_supramarginal_thickness | Correlation | 0.17 | 0.221 | 0.038 |
| | Significance (2-tailed) | 0.085 | 0.025 | 0.704 |
| | df | 101 | 101 | 103 |
| lh_frontalpole_thickness | Correlation | 0.006 | 0.031 | -0.032 |
| | Significance (2-tailed) | 0.946 | 0.701 | 0.69 |
| | df | 151 | 152 | 156 |
| lh_temporalpole_thickness | Correlation | 0.01 | -0.132 | 0.165 |
| | Significance (2-tailed) | 0.901 | 0.109 | 0.041 |
| | df | 147 | 148 | 152 |
| lh_transversetemporal_thickness | Correlation | 0.006 | 0.081 | -0.093 |
| | Significance (2-tailed) | 0.943 | 0.318 | 0.244 |
| | df | 152 | 153 | 157 |
| lh_insula_thickness | Correlation | -0.017 | -0.002 | -0.025 |
| | Significance (2-tailed) | 0.851 | 0.986 | 0.775 |
| | df | 122 | 123 | 127 |
| rh_bankssts_thickness | Correlation | -0.047 | -0.065 | 0.024 |
| | Significance (2-tailed) | 0.58 | 0.446 | 0.78 |
| | df | 137 | 138 | 142 |
| rh_caudalanteriorcingulate_thickness | Correlation | -0.096 | -0.1 | -0.043 |
| | Significance (2-tailed) | 0.271 | 0.246 | 0.615 |
| | df | 132 | 133 | 137 |
| rh_caudalmiddlefrontal_thickness | Correlation | 0.135 | 0.095 | 0.099 |
| | Significance (2-tailed) | 0.103 | 0.252 | 0.227 |
| | df | 144 | 145 | 149 |

| | | | | |
|---|---|---|---|---|
| rh_cuneus_thickness | Correlation | 0.125 | 0.125 | 0.069 |
| | Significance (2-tailed) | 0.178 | 0.174 | 0.447 |
| | df | 116 | 117 | 121 |
| rh_entorhinal_thickness | Correlation | 0.06 | 0.001 | 0.089 |
| | Significance (2-tailed) | 0.46 | 0.987 | 0.266 |
| | df | 151 | 152 | 156 |
| rh_fusiform_thickness | Correlation | -0.047 | -0.032 | -0.034 |
| | Significance (2-tailed) | 0.563 | 0.689 | 0.669 |
| | df | 151 | 152 | 156 |
| rh_inferiorparietal_thickness | Correlation | 0.09 | 0.1 | 0.043 |
| | Significance (2-tailed) | 0.324 | 0.272 | 0.633 |
| | df | 121 | 121 | 126 |
| rh_inferiortemporal_thickness | Correlation | 0.04 | 0.192 | -0.146 |
| | Significance (2-tailed) | 0.668 | 0.04 | 0.114 |
| | df | 113 | 113 | 117 |
| rh_isthmuscingulate_thickness | Correlation | -0.068 | -0.095 | -0.001 |
| | Significance (2-tailed) | 0.401 | 0.241 | 0.987 |
| | df | 152 | 153 | 157 |
| rh_lateraloccipital_thickness | Correlation | 0.127 | 0.126 | 0.052 |
| | Significance (2-tailed) | 0.136 | 0.138 | 0.534 |
| | df | 138 | 138 | 142 |
| rh_lateralorbitofrontal_thickness | Correlation | -0.127 | -0.152 | -0.017 |
| | Significance (2-tailed) | 0.131 | 0.068 | 0.837 |
| | df | 142 | 142 | 146 |
| rh_lingual_thickness | Correlation | 0.097 | 0.059 | 0.097 |
| | Significance (2-tailed) | 0.291 | 0.518 | 0.28 |
| | df | 118 | 119 | 123 |
| rh_medialorbitofrontal_thickness | Correlation | 0.049 | 0.017 | 0.083 |
| | Significance (2-tailed) | 0.615 | 0.861 | 0.384 |
| | df | 108 | 108 | 111 |
| rh_middletemporal_thickness | Correlation | 0.047 | 0.078 | 0 |
| | Significance (2-tailed) | 0.661 | 0.461 | 1 |
| | df | 88 | 89 | 92 |
| rh_parahippocampal_thickness | Correlation | 0.165 | 0.084 | 0.197 |
| | Significance (2-tailed) | 0.041 | 0.3 | 0.013 |
| | df | 152 | 153 | 157 |
| rh_paracentral_thickness | Correlation | 0.092 | 0.126 | -0.006 |
| | Significance (2-tailed) | 0.268 | 0.127 | 0.942 |
| | df | 145 | 146 | 149 |
| rh_parsopercularis_thickness | Correlation | 0.03 | 0.064 | -0.011 |
| | Significance (2-tailed) | 0.72 | 0.433 | 0.896 |
| | df | 147 | 148 | 151 |
| rh_parsorbitalis_thickness | Correlation | 0.027 | -0.047 | 0.132 |
| | Significance (2-tailed) | 0.739 | 0.566 | 0.099 |
| | df | 150 | 151 | 155 |
| rh_parstriangularis_thickness | Correlation | 0.019 | 0.039 | 0.017 |
| | Significance (2-tailed) | 0.815 | 0.637 | 0.833 |

| | | | | |
|---|---|---|---|---|
| | df | 149 | 150 | 154 |
| rh_pericalcarine_thickness | Correlation | 0.204 | 0.096 | 0.192 |
| | Significance (2-tailed) | 0.031 | 0.31 | 0.038 |
| | df | 110 | 111 | 115 |
| rh_postcentral_thickness | Correlation | 0.142 | 0.179 | 0.01 |
| | Significance (2-tailed) | 0.132 | 0.056 | 0.912 |
| | df | 112 | 113 | 115 |
| rh_posteriorcingulate_thickness | Correlation | -0.072 | -0.069 | -0.02 |
| | Significance (2-tailed) | 0.38 | 0.4 | 0.804 |
| | df | 149 | 150 | 154 |
| rh_precentral_thickness | Correlation | 0.172 | 0.181 | 0.079 |
| | Significance (2-tailed) | 0.057 | 0.045 | 0.379 |
| | df | 121 | 122 | 123 |
| rh_precuneus_thickness | Correlation | 0.042 | 0.058 | 0.002 |
| | Significance (2-tailed) | 0.613 | 0.482 | 0.981 |
| | df | 147 | 148 | 152 |
| rh_rostralanteriorcingulate_thickness | Correlation | 0.042 | 0.03 | 0.047 |
| | Significance (2-tailed) | 0.611 | 0.717 | 0.561 |
| | df | 149 | 150 | 154 |
| rh_rostralmiddlefrontal_thickness | Correlation | 0.058 | 0.027 | 0.064 |
| | Significance (2-tailed) | 0.479 | 0.742 | 0.428 |
| | df | 147 | 148 | 152 |
| rh_superiorfrontal_thickness | Correlation | 0.18 | 0.139 | 0.153 |
| | Significance (2-tailed) | 0.044 | 0.118 | 0.081 |
| | df | 124 | 125 | 129 |
| rh_superiorparietal_thickness | Correlation | 0.257 | 0.228 | 0.141 |
| | Significance (2-tailed) | 0.006 | 0.014 | 0.129 |
| | df | 112 | 113 | 116 |
| rh_superiortemporal_thickness | Correlation | 0.027 | 0.028 | 0.029 |
| | Significance (2-tailed) | 0.783 | 0.771 | 0.757 |
| | df | 106 | 106 | 111 |
| rh_supramarginal_thickness | Correlation | 0.13 | 0.144 | 0.075 |
| | Significance (2-tailed) | 0.177 | 0.134 | 0.425 |
| | df | 108 | 108 | 113 |
| rh_frontalpole_thickness | Correlation | -0.016 | -0.06 | 0.035 |
| | Significance (2-tailed) | 0.841 | 0.457 | 0.66 |
| | df | 151 | 152 | 156 |
| rh_temporalpole_thickness | Correlation | 0.011 | 0.038 | -0.025 |
| | Significance (2-tailed) | 0.893 | 0.641 | 0.758 |
| | df | 148 | 149 | 153 |
| rh_transversetemporal_thickness | Correlation | -0.056 | -0.029 | -0.043 |
| | Significance (2-tailed) | 0.492 | 0.723 | 0.592 |
| | df | 152 | 153 | 157 |
| rh_insula_thickness | Correlation | -0.084 | -0.078 | -0.047 |
| | Significance (2-tailed) | 0.363 | 0.395 | 0.604 |
| | df | 117 | 118 | 121 |
| lh_MeanThickness_t | Correlation | 0.081 | 0.093 | 0.025 |

| | | | | |
|---|---|---|---|---|
| hickness | Significance (2-tailed) | 0.319 | 0.248 | 0.755 |
| | df | 152 | 153 | 157 |
| rh_MeanThickness_t | Correlation | 0.115 | 0.094 | 0.083 |
| hickness | Significance (2-tailed) | 0.155 | 0.243 | 0.3 |
| | df | 152 | 153 | 157 |
